# Simple, universal rules predict trophic interaction strengths

**DOI:** 10.1101/2024.07.26.605380

**Authors:** Kyle E. Coblentz, Mark Novak, John P. DeLong

## Abstract

Many drivers of ecological systems exhibit regular scaling relationships, yet the mechanisms explaining these relationships are often unknown. Trophic interaction strengths are no exception, exhibiting scaling relationships with predator and prey traits that lack evolutionary explanations. We propose two rules to explain the scaling of trophic interaction strengths through the relationship between a predator’s feeding rate and its prey’s density -- the so-called predator functional response. First, functional responses allow predators to meet their energetic demands when prey are rare. Second, functional responses approach their maxima near the highest prey densities predators experience. We show that equations derived from these rules predict functional response parameters across over 2,100 functional response experiments and make additional predictions such as their allometric scaling. The two rules thereby offer a potential ultimate explanation for the determinants of trophic interaction strengths, revealing ecologically realized constraints to the complex, adaptive nature of functional response evolution.

## Introduction

Understanding the determinants of predator-prey interaction strengths is key to understanding the stability, diversity, and dynamics of ecosystems (De Ruiter *et al*. 1995; McCann *et al*. 1998; Paine 1992; Yodzis 1981). A fundamental component of trophic interaction strengths is the predator’s functional response that describes how predator feeding rates change with prey density (Holling 1959; Solomon 1949).

Thousands of experiments have measured functional responses for taxa ranging from microbes to large carnivores from nearly every biome on earth (Uiterwaal *et al*. 2022) uncovering a number of variables influencing the underlying parameters (Coblentz *et al*. 2023; DeLong 2021; Pawar *et al*. 2012; Uiterwaal & DeLong 2020; Vucic-Pestic *et al*. 2010). Principal among these variables are predator and prey body sizes whose allometric relationships with functional response parameters serve as the basis for most dynamical food web models (Brose *et al*. 2006; Otto *et al*. 2007; Yodzis & Innes 1992). However, despite the existence and utility of such statistical relationships between functional response parameters and many traits, it remains unclear why these relationships exist as they do.

Early theory suggested that predator biomass consumption should scale to the ¾ power because organisms’ metabolic rates scale to approximately the ¾ power with their masses (Yodzis & Innes 1992). More recent theory considering proximate mechanisms for the strengths of trophic interactions, such as the dimensionality of predator-prey interactions and the allometric scaling of predator and prey movement velocities, has also seen success in predicting certain functional response parameters (i.e. the space clearance/attack/search rate) and their scaling relationships (Pawar *et al*. 2012; Portalier *et al*. 2022). But other parameters (i.e. the asymptotic/maximum feeding rate or its reciprocal, handling time) remain poorly predicted, such that an understanding of the evolutionary drivers of functional responses remains missing. An encompassing means to predict predator foraging rates through theory reflecting their ultimate causes is therefore of significant interest.

We aim to provide such a theory derived from two simple rules for predator foraging. We also examine the ability of our theory to predict functional response parameters and assess several additional theoretical predictions.

### Two Rules for Functional Responses

We posit that predator functional responses meet two conditions. First, a predator’s feeding rate meets its energetic demands at the low prey densities it is likely to experience. This must be true for the long-term persistence of the predator population. Although cases exist in which predator mortality or biomass loss exceeds energy intake and reproduction, as in the declining phases of predator-prey cycles, predator energy intake must balance energetic demand over the long term to avoid extirpation (McCann 2011). Second, the prey densities at which a predator’s feeding rate saturates (i.e. approaches its maximum) should be near the highest prey densities it is likely to experience. This is because, first, feeding rates must saturate at some prey density (Holling 1959; Jeschke *et al*. 2004). Second, if feeding rates saturated at a lower prey density, predators would pay an opportunity cost for missing out on higher feeding rates at the high prey densities they experience. Third, lower handling times leading to higher, less saturated feeding rates at high prey densities: i) have diminishing fitness returns since the highest prey densities are rarely experienced, ii) imply less energy extracted per prey consumed (Okuyama 2010), and iii) require larger ingestive or digestive capacities that are energetically costly (Armstrong & Schindler 2011; McWilliams & Karasov 2001; Secor *et al*. 1994). Although predators may rarely experience these high prey densities, selection on predators may still be strong because taking advantage of these periods of high prey abundance can have outsized effects on predator fitness. For example, Armstrong and Schindler (2011) showed that many fishes have excess digestive capacities to achieve feeding rates up to three times their average rate despite the high cost of maintaining that capacity.

Past studies on functional response evolution have largely focused on relatively simple optimization of predator feeding rates (Abrams 1982; Amarasekare 2022; DeLong & Coblentz 2022). The two rules we posit take a different view in which functional responses are the outcome of the long-term evolution of the traits determining functional responses that balance a multitude of selective pressures. The outcome of this evolutionary process is that predators are minimally capable of persisting when prey are scarce while also taking advantage of occasionally abundant prey.

We derive two equations encapsulating these constraints. The first rule that predators must satisfy their energetic demand at low prey densities can be formalized as

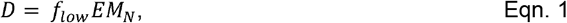

where *D* is predator energetic demand in kJ/day. The rate at which prey are consumed when rare is given by its feeding rate *f*_*low*_, which should be well approximated by a linear functional response at low prey densities (i.e. *f*_*low*_ = *aN*_*low*_, where *a* is the space clearance rate and *N*_*low*_ the low prey density) (Coblentz *et al*. 2023; Novak *et al*. 2024). *E* is the energy density of the prey in kJ/g, and *M*_*N*_ is the mass of the prey in grams to convert the number of prey eaten to the mass of prey eaten. Although we do not explicitly account for lost or otherwise unassimilated prey energy due to undigestible parts, feces, urine, etc. (Yodzis & Innes 1992), these processes could easily be included in eqn. 1 by adjusting the prey energy density or mass.

The second rule that a predator’s feeding rate should saturate near the highest prey densities it experiences can be formalized as

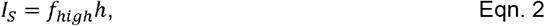

where *I*_*s*_ is the degree of saturation (ranging between 0 and 1) and *h* is the handling time (i.e. the reciprocal of the maximum feeding rate) (Coblentz *et al*. 2023). Assuming a Holling Type II functional response, 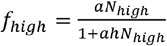 where *N*_*high*_ is the high density of prey likely to be experienced by the predator. *I*_*s*_ also may be interpreted as the fraction of time a predator individual is busy handling prey, or as the fraction of individuals in a predator population handling prey at any given point in time (Supplementary Information) (Coblentz *et al*. 2021; Novak *et al*. 2017). We focus on the Holling Type-II version of the functional response over the Michaelis-Menten form (i.e. 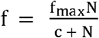 where *f* is the feeding rate, *f*_*max*_ is the maximum feeding rate, and *c* is the half-saturation constant or the prey density at which 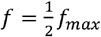). We do so for two reasons. First, the Holling Type II functional response parameters have clear mechanistic interpretations: *a* is the space that would be cleared by a predator in the absence of a time cost to handling prey, and *h* is the time taken away from searching for prey once a prey is captured. Second, the parameters of the Holling Type II functional response have practical interpretations related to feeding rates at high and low prey densities: *a* is the slope of the functional response at the origin, and 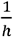 is the maximum feeding rate.

Given equations 1 and 2, we solve for the space clearance rate and handling time as

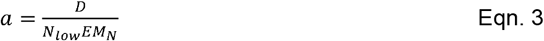

and

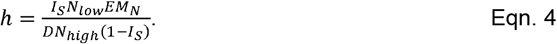

#### Auxiliary Predictions

Beyond providing equations for the space clearance rate and handling time parameters, equations 1 and 2 also make auxiliary predictions. Here, we focus on three of these related to i) the degree of feeding rate saturation across prey densities experienced by predators, ii) the relationship between space clearance rates and handling times, and iii) the allometric scaling of space clearance rates and handling times.

i. Regarding the saturation of predator feeding rates with prey densities, we assume that the functional response is unsaturated at low prey densities and approaches saturation at the high prey densities experienced by predators (eqns. 1 and 2). Given these two assumptions, we also can assess the prey density at which the predator’s feeding rate is half saturated. This occurs when the prey density is equal to 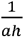 (the half saturation constant in the Michalis-Menten formulation). Using equations 3 and 4, we solve for the half saturation constant as

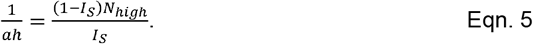 Thus, our theory predicts that, despite different predators experiencing different ranges of prey densities, their degree of saturation across those density ranges is invariant.
ii. Regarding the relationship between space clearance rates and handling times, it follows from eqn. 5 that *a* and *h* are related as

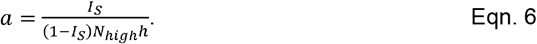

and taking the natural log of both sides,

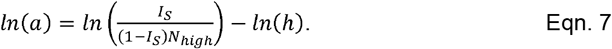 Our theory therefore predicts an inverse (or negative log-log-linear) relationship between space clearance rates and handling times among predators feeding on prey with similar high population sizes.
iii. Last, several of the parameters on the right-hand sides of equations 3 and 4 have well-known scaling relationships with species’ body masses. Specifically, energetic demand defined as predator metabolic rate scales with predator mass (Kleiber’s Law (Brown *et al*. 2004; Kleiber 1932)) and prey densities scale with prey mass (Damuth’s Law (Damuth 1981)). As we can treat the remaining parameters as unrelated to body size, this suggests that the theory outlined here should be capable of predicting the allometric scaling of *a* and *h*.

Below, we take advantage of the FoRAGE database of functional response experiments to examine the predictions of our theory (Uiterwaal *et al*. 2022). We first combine FoRAGE with databases on mass-abundance and metabolic scaling relationships to predict functional response parameters across most studies in FoRAGE. Then, for a subset of FoRAGE studies performed in the field, we examine our theory’s ability to make predictions using system-specific information. Last, we examine the evidence for the auxiliary predictions made by our theory.

## Methods

### Databases, Data Handling, and Predictions

We evaluate the ability of our theory to predict functional response parameters by comparing measured functional response parameters to predictions derived using estimates of the parameters on the right-hand sides of the equations. To do so, we bring together two databases: the FoRAGE compilation from Uiterwaal *et al*. (2022) and a data compilation of eukaryotic species’ masses, metabolic rates, and population densities from Hatton et al. (2019). FoRAGE contains functional response parameter estimates from 3,013 functional response experiments along with additional information, including predator and prey body masses, for most studies. Thus, FoRAGE provides the measured space clearance rates and handling times and values for prey mass in wet weight (*M*_*N*_). To predict space clearance rates and handling times using equations 3 and 4, we still require estimates of energetic demand (*D*), prey energy density (*E*), high and low prey densities (*N*_*low*_ and *N*_*high*_) and the degree of feeding rate saturation at high prey densities (*I*_*s*_).

To obtain estimates of low and high prey densities likely to be experienced by predators (*N*_*low*_ and *N*_*high*_), and predator energy demand (*D*), we combined FoRAGE with mass scaling data from Hatton et al. (2019). To estimate *N*_*low*_ and *N*_*high*_, we used the data from Hatton et al. (2019) containing species’ masses and their measured densities in aerial m^2^. These density estimates include density estimates from the same species in different places or times, as well as estimates across different species. We assume that the variation in the density data reflects both variation in prey densities a predator might experience over space and time and among rare and common species with similar masses. To estimate *N*_*low*_ and *N*_*high*_, for the prey in FoRAGE, we performed Bayesian log-log regressions of density on mass separately for prokaryotes (n = 635), protists (n = 301), invertebrates (n = 778), ectotherm vertebrates (n = 404), mammals (n = 2,852), and birds (n = 603). Using the prey classifications and masses in FoRAGE, we then estimated the posterior predictive distribution for each study-specific prey and extracted the 10^th^ and 90^th^ percentiles as estimates of *N*_*low*_ and *N*_*high*_ respectively. To estimate *D*, we performed Bayesian log-log regressions of basal metabolic rate in kJ/d on predator mass in g separately for protists (n = 365), invertebrates (n = 4,559), ectotherm vertebrates (n = 616), mammals (n = 1,059), and birds (n = 492). We then used the median predicted metabolic rate of each predator in FoRAGE based on its classification and mass as our estimate of *D*. For the Bayesian regressions, we used flat priors on the intercept and slope, and a Student-t prior with degrees of freedom = 3, location = 0, and scale = 2.5 on the standard deviation. We approximated the posterior distribution using 1000 samples each from 4 Markov chains after a 1000 sample warmup. We performed the regressions using the R package ‘brms’ (Bürkner 2017) using Stan (Stan Development Team 2024) through the backend cmdstanr (Gabry *et al*. 2024).

With estimates of *N*_*low*_, *N*_*high*_, and *D*, we then needed estimates of *E* and *I*_*s*_. We assume that these values are approximately constant, setting *E* = 5.6kJ/g wet weight following Brown et al. (2018) and *I*_*S*_ =0.9.Our results are not sensitive to reasonable choices of values for *I*_*S*_ or for the percentiles used for *N*_*low*_ and *N*_*high*_ (see Supplemental Information).

Although the full FoRAGE database contains 3,013 experiments, we restricted our predictions to:

1. Studies including living prey that were not eggs. We did not include studies using eggs as prey as it is unclear whether mass scaling relationships apply to eggs.
2. Studies with handling time estimates greater than 1×10^-6^ days. We did so following previous studies suggesting that studies within FoRAGE with lower values than this cutoff are those for which Type II functional responses have poor fits (Coblentz *et al*. 2023; Kiørboe & Thomas 2020; Uiterwaal & DeLong 2020).
3. Studies with both predator and prey masses. We excluded studies without this information because it is what allows a link between the FoRAGE database and empirical mass scaling relationships.

After excluding the studies that did not meet these criteria, we were left with 2,162 functional response measurements.

Combining the estimates of *N*_*low*_, *N*_*high*_, *D, E*, and *I*_*S*_, we predicted the space clearance rates and handling times for each predator-prey pair in the reduced version of FoRAGE. We compared these predictions to observed values using reduced major axis regression with the R package ‘lmodel2’ (Legendre 2018).

### Field-study Specific Predictions

Forty of the 2,162 FoRAGE experiments in our reduced dataset were performed in the field. Because these field studies include high and low abundances of prey observed during the studies, they provide an opportunity to test functional response parameter predictions using system-specific information rather than mass scaling relationships. To avoid circularity, we excluded studies that estimated predator kill rates using the proportion of predator diets consisting of a prey type coupled with the average mass of that prey and the predator’s daily energetic demand to estimate the number of prey consumed per day. For the remaining studies, we performed a literature search to find information on predator energetic demand (*D*) and prey energy density (*E*). For energy demand, we used estimates of daily energy expenditure, if available. Otherwise, we used basal metabolic rate. For energy density, we used species-specific estimates of kJ/g if they were available. Otherwise, we used studies of prey body composition that gave the percent of prey body mass composed of protein and fat. We used conversions of 16.74 kJ/g for protein and 33.47 kJ/g of fat to calculate prey energy density (Chizzolini *et al*. 1999; Menzies *et al*. 2022). We were able to find the requisite information for seven studies of six mammalian predator-prey pairs. For these studies, we assumed the highest and lowest recorded prey densities as estimates of *N*_*low*_ and *N*_*high*_. Details and references for the estimates are given in the Supplemental Information. As the estimate of mass, we used the mass in FoRAGE (Uiterwaal *et al*. 2022). FoRAGE presents the mass reported in the original study where possible, but if the mass was not available from the original study, the authors of FoRAGE took a stepwise process to estimate masses, for example using length-mass relationships or data from sources other than the original study (for details see (Uiterwaal *et al*. 2022)).

### Auxiliary Predictions

i. To examine whether our theory’s predicted relationship between the predator’s half-saturation constant and prey densities held, we used the values of the space clearance rates and handling times from FoRAGE and the mass-abundance scaling prediction for high prey abundances. To assess the accuracy of the predictions, we calculated the correlation and performed a major axis regression on the observed vs. predicted half saturation constants.
ii. To examine our theory’s prediction of the relationship between space clearance rates and handling times given *I*_*S*_ and the high density of the prey, we used the values of the space clearance rates and handling times from FoRAGE and the mass-abundance scaling predictions of the high prey abundances. To assess the accuracy of the prediction of the relationship between space clearance rates and handling times, we calculated the correlation and performed a major axis regression on the observed vs. predicted space clearance rates.
iii. To assess whether our theory could predict the allometric scaling of space clearance rates and handling times, we derived equations for the scaling of space clearance rates and handling times assuming that predator energy demand (*D*) has a power-law scaling with predator mass, that low and high prey densities (*N*_*low*_ and *N*_*high*_) scale with prey mass, and that predator and prey masses scale with one another. We assumed the other variables are constants unrelated to predator or prey mass. Derivations of these equations are given in the Supplemental Information. We then used log-log least squares regressions to test our predictions, obtaining estimates of the empirical relationships within FoRAGE across the different classifications used in the mass-abundance and mass-metabolism regressions between prey masses and abundances, predator masses and energy demand, and predator and prey masses (see *Databases, Data Handling, and Predictions*).

All analyses were performed in R using v. 4.3.1 (R Core Team 2023)

## Results

### Prediction of a and h

The two rules performed well in predicting empirically estimated space clearance rate and handling time values across their respective 19 and 8 orders of magnitude in variation (Figure 1; log space clearance rate correlation coefficient = 0.82, R^2^ of 1:1 line = 0.81; log handling time correlation coefficient = 0.57, R^2^ of 1:1 line = 0.75). The major axis regression slopes between observed and predicted space clearance rates (*ß* = 0.85, 95% Confidence Interval (CI) = 0.82, 0.87) and handling times (*ß* = 1.14, 95% CI = 1.08, 1.22) suggest a tendency for the slight overestimation of space clearance rates and underestimation of handling times at low values, and the underestimation of space clearance rates and overestimation of handling times at high values (Figure 1).

**Figure 1.**
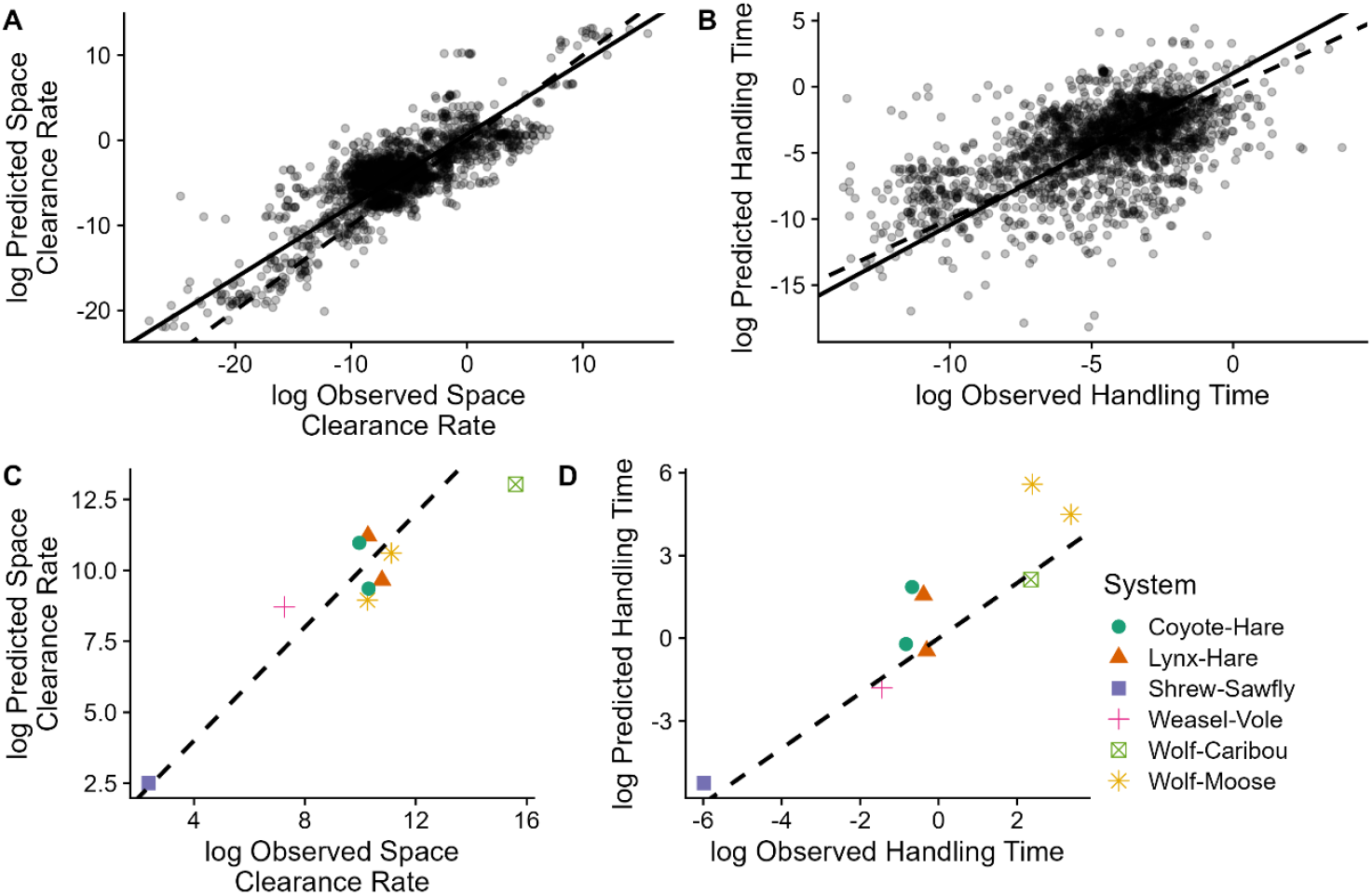
Two simple rules derived from predator feeding rates at low and high prey densities predict space clearance rates and handling times using either generic estimates from empirical mass scaling relationships (A,B) or system-specific estimates from field studies (C,D). Dashed lines are 1:1 lines and solid lines are major axis regressions. See the Supplemental Information for versions of panels A and B with color-coded predator and prey body size information.

Alternative means for determining low and high prey densities and levels of predator saturation shift the points in Figure 1, but do not influence the predictive power of the two rules (Supplementary Information). Indeed, the tendency for over/under-estimation likely reflects insightful across-system variation not reflected in estimates from mass-scaling relationships (see below).

We also predicted the functional response parameters for the subset of the studies that were field studies. For the seven studies of mammalian predators, we again find that the predictions fall near the 1:1 line but without the same systematic tendency for over/under-estimation (Figure 1C-D).

### Auxiliary Predictions

i. The relationship between the empirical half-saturation constant values and high prey densities, modified by the degree of saturation, supports our theory’s prediction (correlation = 0.86, slope of major axis regression (95% CI) = 0.88 (0.86,0.9), Fig. 2A; see the Supplemental Information for a sensitivity analysis).
ii. Space clearance rates and handling times were inversely related to each other when accounting for the high prey densities (Figure 2B,C). Without accounting for the high prey densities, there is little apparent relationship between space clearance rates and handling times (Figure 2B). However, once the high prey densities are accounted for, the negative relationship between space clearance rates and handling times is evident (see the color coding in Figure 2B). Examining the prediction of the relationship between handling times modified by high prey densities and the degree of predator feeding rate saturation at high prey densities supports our theory’s prediction, albeit with a tendency to underpredict the space clearance rates (correlation = 0.84, intercept of major axis regression (95% CI) = -1.5 (-1.6,-1.4), slope of major axis regression (95%CI) = 1.07 (1.04,1.1)).
iii. Space clearance rates and handling times scale with predator and prey body masses as predicted. Specifically, the empirical scaling of the parameters we consider as being related to body size (those besides *I*_*S*_ and *E*) are 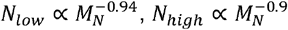 and 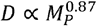 where *M*_*P*_ is the mass of the predator and the other parameters are defined above. Given these scaling relationships and the first rule, it follows that the allometric relationship for the space clearance rate should be 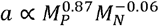.Given that in the dataset 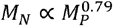,we therefore predict that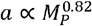. This prediction matches the observed scaling of 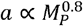 (95% CI = 0.78, 0.83; Figure 3A). Similarly, the allometric relationship for handling time should be 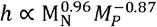.Thus, we predict 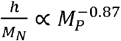.This prediction also matches the observed scaling of 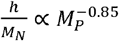 (95% CI = -0.87, -0.82; Figure 3B; See Supplemental Information for derivations of scaling relationships).

**Figure 2.**
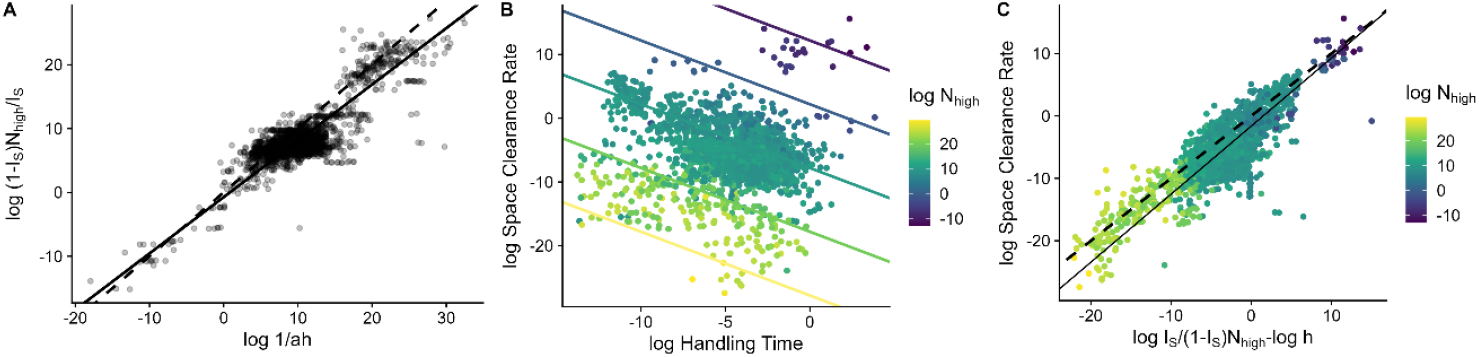
The two rules predict empirical estimates of the half-saturation constant (*1/ah*) (A), as well as the relationship between space clearance rates and handling times (B,C) using only knowledge of the saturation experienced at high experienced prey densities and those high prey densities. The dashed lines in A and C are the 1:1 lines and the solid lines are major axis regressions. The lines in B are the predicted relationships between space clearance rates and handling times for different values of *N*_*high*_. See the Supplemental Information for versions of panels A with color-coded predator and prey body size information.

**Figure 3.**
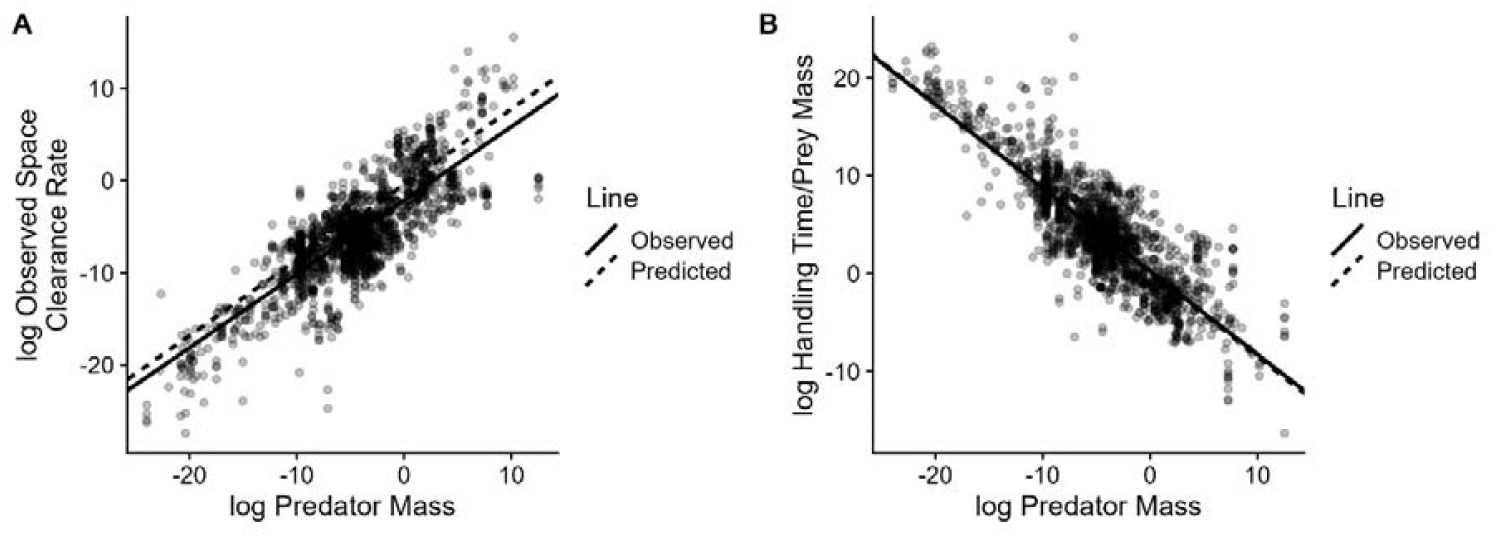
Functional response rules and empirical scaling relationships between parameters and prey and predator masses predict that the log space clearance rate increases with log predator mass with a slope of 0.82 (observed empirical estimate: 0.8, A) and that the log of the prey mass-specific handling time (handling time/prey mass) decreases with log consumer mass with a slope of -0.87 (observed empirical estimate: -0.86; B). See the Supplemental Information for versions of the panels with color-coded predator and prey body size information.

## Discussion

We demonstrate that two simple rules predict the parameters of predator functional responses and their scaling relationships. In contrast to prior work, our theory provides an ultimate explanation for the strengths of trophic interactions rather than focusing on proximate mechanisms. Our approach generates predictions of space clearance rates of predators with similar accuracy as prior theory (i.e. most predictions within ∼ 3 orders of magnitude of the observed values; Pawar *et al*. 2012; Portalier *et al*. 2022), but also predicts handling times, which have been predicted poorly by previous work (Portalier *et al*. 2022). In addition, our theory offers two practical advantages. First, it uses parameters that are generally easier to measure than those required by previous theory (e.g. predator movement velocities and prey detection distances). Second, our theory’s foundation in ultimate causes suggests potential applications beyond environmental conditions and taxa in which trophic interaction strengths have been measured, addressing a limitation of current statistical approaches to prediction (DiFiore & Stier 2023).

Our two rules emerge from a view of functional responses as products of a long-term eco-co-evolutionary process involving predators, their prey, and the environment including the effects of other species and their interactions (DeLong 2020; Gutiérrez Al-Khudhairy & Rossberg 2022; Wickman *et al*. 2024). A predator’s fitness depends not only on its ability to procure food; it also depends on its ability to avoid its own predators, find mates, etc. (Jeschke 2007). Therefore, we suggest that eco-evolutionary processes combine such that trophic interaction strengths reflect the ability of a predator to minimally meet energetic demands during busts of low prey densities while also taking advantage of instances of booms of high prey densities. This contrasts with previous work focusing on the optimization of space clearance rates and handling times to maximize energy intake. This view also may help explain why consumers tend not to overexploit their prey (Gutiérrez Al-Khudhairy & Rossberg 2022; Vuorinen *et al*. 2021).

As many complex behaviors and phenotypes are the product of multiple traits under multiple selective pressures, our approach of focusing on functional outcomes rather than specific processes may offer a way forward for understanding and predicting other complex phenotypes such as thermal responses and competitive abilities and their resulting ecological scaling relationships.

The theory developed here also makes predictions beyond the values of space clearance rates and handling times. The first is the prediction that predators show consistent patterns in how their feeding rates saturate across prey densities: feeding rates are largely unsaturated at low prey densities, reach half-saturation at an intermediate fraction of the highest prey densities, and only become fully saturated at the highest prey densities encountered. This result is consistent with that of Coblentz et al. (2023) who found that predator feeding rates were generally unsaturated at the typical prey densities predators are likely to experience. Indeed, given the highly skewed frequencies of species abundances spatially and temporally in nature (Brown *et al*. 1995; Halley & Inchausti 2002), the predicted prey density at which half-saturation occurs within our data is typically 2.4 times higher (standard deviation = 0.4) than the predicted median prey density. This suggests that predators only rarely feed near their maximum feeding rates, implying that predation pressure should dynamically change with changes in prey densities.

The second auxiliary prediction from our theory is a negative relationship between space clearance rates and handling times, a pattern previously found by Kiørboe & Thomas (2020). These authors interpreted this relationship as a slow-fast continuum in trophic interactions that appeared to contradict the gleaner-opportunist tradeoff – a fundamental concept in ecology that helps explain species coexistence under variable resources (Armstrong & McGehee 1980; Grover 1990). Under the gleaner-opportunist tradeoff, ‘gleaners’ perform better at low resource levels while ‘opportunists’ take advantage of high resource situations. If ingestion rates directly determine growth rates, the gleaner-opportunist tradeoff would predict a positive relationship between space clearance rates and handling times (Kiørboe & Thomas 2020), a pattern that is opposite to that which we and Kiørboe & Thomas (2020) observed. Our results provide a potential resolution to this apparent contradiction by considering predator energy demand. Our theory suggests that the negative relationship between space clearance rates and handling times for predators consuming similarly sized prey occurs due to differences in energy demand -- energy demand appears in the numerator and denominator of the equations for space clearance rates and handling times respectively. Thus, predators with higher energy demands exhibit higher space clearance rates and lower handling times, whereas predators with lower energy demands exhibit the opposite pattern. Although high-energy-demand predators may consume more prey at low prey availability, they also incur higher energetic costs under these conditions compared to low-energy-demand predators. This energetic tradeoff provides a mechanism for the co-occurrence of a gleaner-opportunist tradeoff with the presence of a slow-fast continuum in functional responses parameter.

The third auxiliary prediction from our theory is the allometric scaling of space clearance rates and handling times that play a central role in dynamical food web models (Brose *et al*. 2006; Otto *et al*. 2007; Yodzis & Innes 1992). Existing allometric scaling relationships of functional response parameters generally fall into two groups. The first are studies following Yodzis and Innes (1992) who argued that predators’ maximum feeding rates should scale with predator mass to the same exponent as metabolic rate and that the half-saturation constant in units of prey mass is independent of prey and predator masses. Translating this scaling to our own, we find that the two approaches predict similar allometric scaling estimates (prey-mass specific scalings of handling times of -0.87 under both approaches and space clearance rate scalings of 0.82 in our theory and 0.87 for Yodzis and Innes (1992); Supplemental Information S6). Although this convergence in predictions is likely due to the central role played by predator metabolic rate scaling, we note that our approach also predicts the pre-factor of the relationship (Figure 3) and provides a potential explanation for the lack of allometric scaling of the half-saturation constant that is left unexplained in Yodzis and Innes (1992). The second group of allometric scaling relationships focuses on the space clearance rate (McGill & Mittelbach 2006; Pawar *et al*. 2012; Rall *et al*. 2012) and derives different scaling relationships between space clearance rates and predator mass depending on whether predators and prey interact in two or three dimensions. These studies predict that space clearance rates scale with predator body mass to an exponent between 0.58-0.68 in two dimensions and 0.92-1.07 in three dimensions. Our predicted and observed scaling exponent of 0.82 falls between these ranges. Unfortunately, we cannot derive dimension-specific allometric scaling relationships using our current data because the dataset from which we derive our abundance estimates (Hatton *et al*. 2019) is in a single dimension (aerial m^2^) and does not have information on the dimension in which organisms live or interact with their predators. Future work incorporating dimension-specific scaling relationships into our theory will thus be useful for integrating the ultimate and proximate mechanisms determining trophic interaction strengths.

The rules we derive also make specific, testable predictions regarding how the environment and global climate change are likely to influence trophic interaction strengths, as many of the parameters describing our rules are sensitive to the environment. For ectotherms, predator energetic demand is expected to increase with increasing temperature (Brown *et al*. 2004), whereas prey population size, body size and potentially energy density are likely to decrease (Amarasekare & Savage 2012; Atkinson 1994; Bernhardt *et al*. 2018; Brett *et al*. 1969; Outhwaite *et al*. 2022). These concurrent changes will alter the functional response parameters required to satisfy our rules, suggesting the potential for maladaptive trophic interactions under climate change.

Changes in parameters also are likely to occur over other environmental gradients, such as productivity gradients (Novak 2013), which may allow for the prediction of changes in top-down interaction strengths and their consequences for ecosystems. Furthermore, although our analyses have generally focused on interspecific differences in trophic interactions, this theory also may be applicable to intraspecific variation in predators and prey to explain intraspecific variation in functional response parameters (Bolnick *et al*. 2011; Coblentz *et al*. 2021).

The success of our theory in predicting functional response parameters across field and laboratory studies opens several promising directions for future theoretical and empirical insights into trophic interaction strengths. First, our theory provides predictions about which trophic interactions within communities are likely to be strong, helping to identify potential keystone interactions that may disproportionately impact communities. Second, although our theory has focused on pairwise trophic interactions, it provides the foundation for a theory on generalist predators to embrace the additional complexity present in food webs in which predators consume multiple prey species. Last, although we have focused specifically on traditional predators, it may be possible to extend the theory to other consumer types such as herbivores, detritivores, and parasites. By deepening our understanding of the ultimate causes governing trophic interaction strengths, this work promises to provide important insights into the functioning and dynamics of ecosystems and their responses to ongoing global change.

## Acknowledgments

KEC and JPD were partially supported by a James S. McDonnel Foundation Studying Complex Systems Scholar grant and a National Science Foundation Organismal Responses to Climate Change (ORCC-2307464) grant to JPD. MN was supported by a National Science Foundation Division of Environmental Biology grant (DEB-2129758).

## References

Abrams, P.A. (1982). Functional Responses of Optimal Foragers. The American Naturalist, 120, 382–390.

Amarasekare, P. (2022). Ecological Constraints on the Evolution of Consumer Functional Responses. Frontiers in Ecology and Evolution, 10.

Amarasekare, P. & Savage, V. (2012). A Framework for Elucidating the Temperature Dependence of Fitness. The American Naturalist, 179, 178–191.

Armstrong, J.B. & Schindler, D.E. (2011). Excess digestive capacity in predators reflects a life of feast and famine. Nature, 476, 84–87.

Armstrong, R.A. & McGehee, R. (1980). Competitive Exclusion. The American Naturalist, 115, 151–170.

Atkinson, D. (1994). Temperature and Organism Size—A Biological Law for Ectotherms? In: Advances in Ecological Research (eds. Begon, M. & Fitter, A.H.). Academic Press, pp. 1–58.

Bernhardt, J.R., Sunday, J.M. & O’Connor, M.I. (2018). Metabolic Theory and the Temperature-Size Rule Explain the Temperature Dependence of Population Carrying Capacity. The American Naturalist, 192, 687–697.

Bolnick, D.I., Amarasekare, P., Araújo, M.S., Bürger, R., Levine, J.M., Novak, M., et al. (2011). Why intraspecific trait variation matters in community ecology. Trends in Ecology & Evolution, 26, 183–192.

Brett, J.R., Shelbourn, J.E. & Shoop, C.T. (1969). Growth Rate and Body Composition of Fingerling Sockeye Salmon, Oncorhynchus nerka, in relation to Temperature and Ration Size. J. Fish. Res. Bd. Can., 26, 2363–2394.

Brose, U., Williams, R.J. & Martinez, N.D. (2006). Allometric scaling enhances stability in complex food webs. Ecology Letters, 9, 1228–1236.

Brown, J.H., Gillooly, J.F., Allen, A.P., Savage, V.M. & West, G.B. (2004). Toward a Metabolic Theory of Ecology. Ecology, 85, 1771–1789.

Brown, J.H., Hall, C.A.S. & Sibly, R.M. (2018). Equal fitness paradigm explained by a trade-off between generation time and energy production rate. Nat Ecol Evol, 2, 262–268.

Brown, J.H., Mehlman, D.W. & Stevens, G.C. (1995). Spatial Variation in Abundance. Ecology, 76, 2028–2043.

Bürkner, P.-C. (2017). brms: An R Package for Bayesian Multilevel Models Using Stan. Journal of Statistical Software, 80, 1–28.

Chizzolini, R., Zanardi, E., Dorigoni, V. & Ghidini, S. (1999). Calorific value and cholesterol content of normal and low-fat meat and meat products. Trends in Food Science & Technology, 10, 119–128.

Coblentz, K.E., Merhoff, S. & Novak, M. (2021). Quantifying the effects of intraspecific variation on predator feeding rates through nonlinear averaging. Functional Ecology, 35, 1560–1571.

Coblentz, K.E., Novak, M. & DeLong, J.P. (2023). Predator feeding rates may often be unsaturated under typical prey densities. Ecology Letters, 26, 302–312.

Damuth, J. (1981). Population density and body size in mammals. Nature, 290, 699–700.

De Ruiter, P.C., Neutel, A.-M. & Moore, J.C. (1995). Energetics, Patterns of Interaction Strengths, and Stability in Real Ecosystems. Science, 269, 1257–1260.

DeLong, J.P. (2020). Detecting the Signature of Body Mass Evolution in the Broad-Scale Architecture of Food Webs. The American Naturalist, 196, 443–453.

DeLong, J.P. (2021). Predator Ecology: Evolutionary Ecology of the Functional Response. Oxford University Press, Oxford, New York.

DeLong, J.P. & Coblentz, K.E. (2022). Prey diversity constrains the adaptive potential of predator foraging traits. Oikos, 2022.

DiFiore, B.P. & Stier, A.C. (2023). Variation in body size drives spatial and temporal variation in lobster–urchin interaction strength. Journal of Animal Ecology, 92, 1075–1088.

Gabry, J., Češnovar, R., Johnson, A. & Bronder, S. (2024). cmdstanr: R Interface to “CmdStan.”

Grover, J.P. (1990). Resource Competition in a Variable Environment: Phytoplankton Growing According to Monod’s Model. The American Naturalist, 136, 771–789.

Gutiérrez Al-Khudhairy, O.U. & Rossberg, A.G. (2022). Evolution of prudent predation in complex food webs. Ecology Letters, 25, 1055–1074.

Halley, J. & Inchausti, P. (2002). Lognormality in Ecological Time Series. Oikos, 99, 518–530.

Hatton, I.A., Dobson, A.P., Storch, D., Galbraith, E.D. & Loreau, M. (2019). Linking scaling laws across eukaryotes. Proceedings of the National Academy of Sciences, 116, 21616–21622.

Holling, C.S. (1959). The Components of Predation as Revealed by a Study of Small-Mammal Predation of the European Pine Sawfly1. The Canadian Entomologist, 91, 293–320.

Jeschke, J.M. (2007). When carnivores are “full and lazy.” Oecologia, 152, 357–364.

Jeschke, J.M., Kopp, M. & Tollrian, R. (2004). Consumer-food systems: why type I functional responses are exclusive to filter feeders. Biological Reviews, 79, 337–349.

Kiørboe, T. & Thomas, M.K. (2020). Heterotrophic eukaryotes show a slow-fast continuum, not a gleaner–exploiter trade-off. Proceedings of the National Academy of Sciences, 117, 24893–24899.

Kleiber, M. (1932). Body size and metabolism. Hilgardia, 6, 315–353.

Legendre, P. (2018). lmodel2: Model II Regression.

McCann, K., Hastings, A. & Huxel, G.R. (1998). Weak trophic interactions and the balance of nature. Nature, 395, 794–798.

McCann, K.S. (2011). Food Webs (MPB-50). Food Webs (MPB-50). Princeton University Press.

McGill, B.J. & Mittelbach, G.G. (2006). An allometric vision and motion model to predict prey encounter rates. Evol Ecol Res, 8, 691–701.

McWilliams, S.R. & Karasov, W.H. (2001). Phenotypic flexibility in digestive system structure and function in migratory birds and its ecological significance. Comparative Biochemistry and Physiology Part A: Molecular & Integrative Physiology, The Physiological Consequences of Feeding in Animals, 128, 577–591.

Menzies, A.K., Studd, E.K., Seguin, J.L., Derbyshire, R.E., Murray, D.L., Boutin, S., et al. (2022). Activity, heart rate, and energy expenditure of a cold-climate mesocarnivore, the Canada lynx (Lynx canadensis). Can. J. Zool., 100, 261–272.

Novak, M. (2013). Trophic omnivory across a productivity gradient: intraguild predation theory and the structure and strength of species interactions. Proceedings of the Royal Society B: Biological Sciences, 280, 20131415.

Novak, M., Coblentz, K.E. & DeLong, J.P. (2024). In defense of the Type I functional response: The frequency and population-dynamic effects of feeding on multiple prey at a time.

Novak, M., Wolf, C., Coblentz, K.E. & Shepard, I.D. (2017). Quantifying predator dependence in the functional response of generalist predators. Ecology Letters, 20, 761–769.

Okuyama, T. (2010). Prey density-dependent handling time in a predator-prey model. Community Ecology, 11, 91–96.

Otto, S.B., Rall, B.C. & Brose, U. (2007). Allometric degree distributions facilitate food-web stability. Nature, 450, 1226–1229.

Outhwaite, C.L., McCann, P. & Newbold, T. (2022). Agriculture and climate change are reshaping insect biodiversity worldwide. Nature, 605, 97–102.

Paine, R.T. (1992). Food-web analysis through field measurement of per capita interaction strength. Nature, 355, 73–75.

Pawar, S., Dell, A.I., & Van M. Savage. (2012). Dimensionality of consumer search space drives trophic interaction strengths. Nature, 486, 485–489.

Portalier, S.M.J., Fussmann, G.F., Loreau, M. & Cherif, M. (2022). Inferring Size-Based Functional Responses From the Physical Properties of the Medium. Front. Ecol. Evol., 9.

R Core Team. (2023). R: A Language and Environment for Statistical Computing. R Foundation for Statistical Computing, Vienna, Austria.

Rall, B.C., Brose, U., Hartvig, M., Kalinkat, G., Schwarzmüller, F., Vucic-Pestic, O., et al. (2012). Universal temperature and body-mass scaling of feeding rates. Phil. Trans. R. Soc. B, 367, 2923–2934.

Secor, S.M., Stein, E.D. & Diamond, J. (1994). Rapid upregulation of snake intestine in response to feeding: a new model of intestinal adaptation. American Journal of Physiology-Gastrointestinal and Liver Physiology, 266, G695–G705.

Solomon, M.E. (1949). The Natural Control of Animal Populations. Journal of Animal Ecology, 18, 1–35.

Stan Development Team. (2024). Stan Modeling Language Users Guide and Reference Manual, version 2.35.

Uiterwaal, S.F. & DeLong, J.P. (2020). Functional responses are maximized at intermediate temperatures. Ecology, 101, e02975.

Uiterwaal, S.F., Lagerstrom, I.T., Lyon, S.R. & DeLong, J.P. (2022). FoRAGE database: A compilation of functional responses for consumers and parasitoids. Ecology, 103, e3706.

Vucic-Pestic, O., Rall, B.C., Kalinkat, G. & Brose, U. (2010). Allometric functional response model: body masses constrain interaction strengths. Journal of Animal Ecology, 79, 249–256.

Vuorinen, K.E.M., Oksanen, T., Oksanen, L., Vuorisalo, T. & Speed, J.D.M. (2021). Why don’t all species overexploit? Oikos, 130, 1835–1848.

Wickman, J., Litchman, E. & Klausmeier, C.A. (2024). Eco-evolutionary emergence of macroecological scaling in plankton communities. Science, 383, 777–782.

Yodzis, P. (1981). The stability of real ecosystems. Nature, 289, 674–676.

Yodzis, P. & Innes, S. (1992). Body Size and Consumer-Resource Dynamics. The American Naturalist, 139, 1151–1175.

