## Supplemental Information for "Simple, universal rules predict trophic interaction strengths"

**Contents	Page**

**S1 *I_S_* as the Fraction of Predators Feeding	2**

**S2 Derivation of Allometric Scaling between Predator and Prey Masses and**

**Functional Response Parameters	3**

**S3 Field-study Specific Information and Sources	6**

**S4 Supplemental Figures with Predator and Prey Body Masses	8**

**S5 Sensitivity Analysis of Choices for *I_S_*, *N_low_*, and *N_high_*	10**

**S6 Comparison of Allometric Scaling Predictions with Yodzis and Innes (1992)	24**

**Supplementary References	27**

**S1 *I_S_* as the Fraction of Predators Feeding**

Here we show that the index of predator feeding rate saturation introduced in the main text (*I_S_*) is equivalent to the fraction of predators within a population that are handling prey under steady-state conditions (Coblentz *et al.* 2023; Novak *et al.* 2017).

Assume that a population of predators feed with feeding rate *f* and that the predators spend time *h* handling each prey after it is captured. If a predator forages over time *T*, then the total number of prey eaten by the predator will be *fT* and the total time spent handling prey will be *fTh*. The corresponding fraction of total time that the predator spends feeding is then *fh*. Assuming that predators are identical and independent, the fraction of predators feeding at a given moment in population is then also *fh*. Thus, the fraction of predators within a population that are handling prey under steady state conditions is equivalent to *I_S_* which also equals *fh*. Specifically, with a Holling Type II functional response where $f=\frac{aN}{1+ahN}$, we get that $I_{S}=\frac{ahN}{1+ahN}$ as in Equation 2 of the main text with $N$ being replaced by $N_{high}$.

**S2 Derivation of Allometric Scaling between Predator and Prey Masses and**

**Functional Response Parameters**

Here we derive the predicted allometric scaling relationships between predator and prey masses and the functional response parameters.

For the space clearance rate (*a*) and handling time (*h*), we have that

$a=\frac{D}{N_{low}EM_{N}}$	Eqn. S2.1

and

$h=\frac{I_{S}N_{low}EM_{N}}{\left( 1-I_{S} \right)DN_{high}}$,	Eqn. S2.2

where $D$ is the energy demand of the predator, $N_{low}$ is a low density of prey likely to be experienced by the predator, $E$ is the energy density of the prey, $M_{N}$ is the mass of the prey, and *I_S_* is an index of feeding rate saturation at the $N_{high}$ prey density that the predator is likely experience. In these equations, the prey densities scale with prey body size, predator energy demand scales with predator mass, and predator and prey masses scale with one another. We consider all other parameters to be independent of predator and prey body masses and assume values for them as in the main text.

To simplify our analysis, we express the parameters that scale with predator or prey body masses on a logarithmic scale as follows. (Definitions for each pre-factor and exponent parameter are given in Table S2.1.)

$D=\delta_{0}M_{P}^{\delta}$,

$N_{low}=\eta_{0,low}M_{N}^{\eta_{low}}$,

$N_{high}=\eta_{0,high}M_{N}^{\eta_{high}}$.

The predator and high and low prey densities are related to one another as

$$M_{P}=\mu_{0,N}M_{N}^{\mu_{N}}$$

and

$M_{N}=\mu_{0,P}M_{P}^{\mu_{P}}$.

We next incorporate these scaling relationships into equations S2.1 and S2.2. We begin by substituting the scalings for *D* and *N_low_* into equation S2.1 to obtain

$a=\frac{\delta_{0}M_{P}^{\delta}}{\eta_{0,low}M_{N}^{\eta_{low}}EM_{N}}$.

Separating the terms that do not depend on predator or prey masses then gives

$a=\frac{\delta_{0}}{\eta_{0,low}E}M_{P}^{\delta}M_{N}^{-\eta_{low}-1}$.

Using the scaling relationship between *M_N_* and *M_P_*, we isolate the scaling of the space clearance rate on predator mass to obtain

$a=\frac{\delta_{0}}{\eta_{0,low}E}M_{P}^{\delta}\mu_{0,P}^{-\eta low-1}M_{P}^{\mu_{P}\left( -\eta_{low}-1 \right)}= \frac{\delta_{0}\mu_{0,P}^{-\eta_{low}-1}}{\eta_{0,low}E}M_{P}^{\delta+\mu_{P}\left( -\eta_{low}-1 \right)}$	Eqn. S2.3

For the handling time, we substitute the scalings for *D*, *N_high_*, and *N_low_* into equation S2.2 to obtain

$h=\frac{I_{S}\eta_{0,low}M_{N}^{\eta_{low}}EM_{N}}{\left( 1-I_{S} \right)\delta_{0}M_{P}^{\delta}\eta_{0,high}M_{N}^{\eta_{high}}}$.

Separating the terms that do not depend on predator or prey masses gives

$h=\frac{I_{S}\eta_{0,low}E}{\left( 1-I_{S} \right)\delta_{0}\eta_{0,high}}M_{N}^{1+\eta_{low}-\eta_{high}}M_{P}^{-\delta}$.	Eqn. S2.4

Equations S3.3 and S3.4 predict the relationships of prey and predator body mass and space clearance rate and handling time parameters when these are plotted on the log-log scale. To evaluate these predictions, we assumed *I_S_* = 0.9, *E* = 5.6kJ/g, and used the 90^th^ and 10^th^ percentiles of the empirical posterior predictive distributions for *N_high_* and *N_low_* respectively. All other empirically determined parameters are given in Table S2.1. With these values specified, equation S2.3. reduces to

$$a= 0.654M_{P}^{0.82}$$

while equation S2.4 reduces to

$h=0.11M_{P}^{-0.87}M_{N}^{0.96}$,

the latter of these leading to the approximation

$$\frac{h}{M_{N}}\approx1.11M_{P}^{-0.87}$$

as presented in the main text and plotted in Figure 3.

**Table S2.1.** Parameters defining the allometric scaling relationships between predator and prey masses and the parameters in the equations for space clearance rates and handling times, along with their definitions and their estimated values and 95% confidence intervals (CI) from log-log regressions.

| **Parameter** | **Definition** | **Value**  **(95% CI)** |
| --- | --- | --- |
| $\delta_{0}$ | The pre-factor for the scaling between predator mass and predator metabolic demand | 0.059  (0.058,0.06) |
| $\delta$ | The exponent of the scaling relationship between predator mass and predator metabolic demand | 0.867  (0.864,0.87) |
| $\eta_{0,low}$ | The pre-factor for the scaling relationship between prey mass and low prey densities | 0.022  (0.02,0.025) |
| $\eta_{low}$ | The exponent of the scaling relationship between prey mass and low prey densities | -0.94  (-0.95,-0.93) |
| $\eta_{0,high}$ | The pre-factor for the scaling relationship between prey mass and high prey densities | 17.14  (15.62, 18.82) |
| $\eta_{high}$ | The exponent of the scaling relationship between prey mass and high prey densities | -0.90  (-0.91,-0.89) |
| $\mu_{0,N}$ | The pre-factor of the scaling relationship between prey mass and predator mass | 11.26  (8.64,14.67) |
| $\mu_{N}$ | The exponent of the scaling relationship between prey mass and predator mass | 0.8  (0.77,0.82) |
| $\mu_{0,P}$ | The pre-factor of the scaling relationship between predator mass and prey mass | 0.0055  (0.0046,0.0066) |
| $\mu_{P}$ | The exponent of the scaling relationship between predator mass and prey mass | 0.79  (0.76,0.82) |

**S3 Field-study Specific Information and Sources**

Here we provide more detailed information on the parameters we used to predict space clearance rates and handling times for the subset of field studies that occur in the FoRAGE database (Uiterwaal *et al.* 2022).

We first subset the FoRAGE database to only include the 40 functional response measurements that were from field studies. To avoid potential circularity, we then removed studies that did not directly estimate predator kill rates, but rather estimated kill rates using the proportion of prey making up the predator population’s diet and the daily energetic demand of the predators (e.g. (Korpimaki & Norrdahl 1991; Zalewski *et al.* 1995)). For each study, we then determined whether we could find the requisite information to parameterize the equations for the space clearance rates and handling times, namely, the prey mass and energy density, the energy demand of the predator, and the high and low prey densities.

We assumed that the high and low prey densities presented in the study (converted to m^2^) were representative of high and low densities experienced by the predators. We note that one study of weasels feeding on voles was an experimental field study in outdoor enclosures in which prey densities were determined by the experimenters (Sundell *et al.* 2000). However, the range of densities used in the experiment were informed by the variation in natural vole abundances (Sundell *et al.* 2000).

Prey mass is also already recorded for each prey in FoRAGE. For prey energy density, we first used Google Scholar to search for the species and the terms ‘energy density’ or ‘energy content’. If we could not find a reference for the energy density of the prey this way, we performed a Google Scholar search of the species and ‘body composition’ to find studies reporting the percent of the prey mass composed of protein and fat. We then used conversions for the kJ/g of fat and protein and the mass of the prey to determine the energy density of the prey (Chizzolini *et al.* 1999).

To determine the energy demand of the predator, we searched for information on daily energetic expenditure or metabolic rate of the predator. If we were able to find both, we preferentially used daily energetic expenditure as this is more likely to reflect the energy demand of predators under field conditions.

Last, we assumed that the degree of saturation in the predators at high prey densities (*I_S_*) was 0.9 as we did for the other analyses in the main text.

If we were unable to find any of the requisite information for a study, it was dropped from the analysis. Overall, this led to a total of seven mammalian predators that could be used for the analysis. Below, we provide a table including each of the studies, the estimates for each of the requisite parameters from sources other than FoRAGE, and the references from which we derived the parameter estimates (Table S3.1).

**Table S3.1.** Information used to predict space clearance rates and handling times in field studies.

| Predator | Prey | Predator Energy Demand Type | Predator Energy Demand (kJ/Day) | Prey Energy Density Type | Prey Energy Density (kJ/g) |
| --- | --- | --- | --- | --- | --- |
| *Lynx canadensis* (O’Donoghue *et al.* 1998) | *Lepus americanus* | Daily Energy Expenditure (Menzies *et al.* 2022) | 6048 | From Body Composition (Menzies *et al.* 2022) | 4.5 |
| *Canis latrans* (O’Donoghue *et al.* 1998) | *Lepus americanus* | Daily Energy Expenditure (Carbone *et al.* 2007) | 4511 | From Body Composition (Menzies *et al.* 2022) | 4.5 |
| *Lynx canadensis* (Chan *et al.* 2017) | *Lepus americanus* | Daily Energy Expenditure (Menzies *et al.* 2022) | 6048 | From Body Composition (Menzies *et al.* 2022) | 4.5 |
| *Canis latrans* (Chan *et al.* 2017) | *Lepus americanus* | Daily Energy Expenditure (Carbone *et al.* 2007) | 4511 | From Body Composition (Menzies *et al.* 2022) | 4.5 |
| *Mustela nivalis* (Sundell *et al.* 2000) | *Microtus rossiameridonalis* | Daily Energy Expenditure (Carbone *et al.* 2007) | 215.5 | From Body Composition (Gorecki 1965) | 6.3 |
| *Canis lupus* (Dale *et al.* 1994) | *Rangifer tarandus* | Daily Energy Expenditure (Menzies *et al.* 2022) | 25920 | From Body Composition (Cook *et al.* 2021) | 7.3 |
| *Canis lupus* (Messier 1994) | *Alces alces* | Daily Energy Expenditure (Menzies *et al.* 2022) | 25920 | From Body Composition (Schwartz *et al.* 1988) | 8.1 |
| *Canis lupus* (Vucetich *et al.* 2002) | *Alces alces* | Daily Energy Expenditure (Menzies *et al.* 2022) | 25920 | From Body Composition (Schwartz *et al.* 1988) | 8.1 |
| *Blarina brevicauda* (Holling 1959b) | *Neodiprion sertifer* | Basal Metabolic Rate (Buckner 1964) | 40.59 | Directly Measured (Buckner 1964) | 5 |

**S4 Supplemental Figures with Predator and Prey Body Sizes**

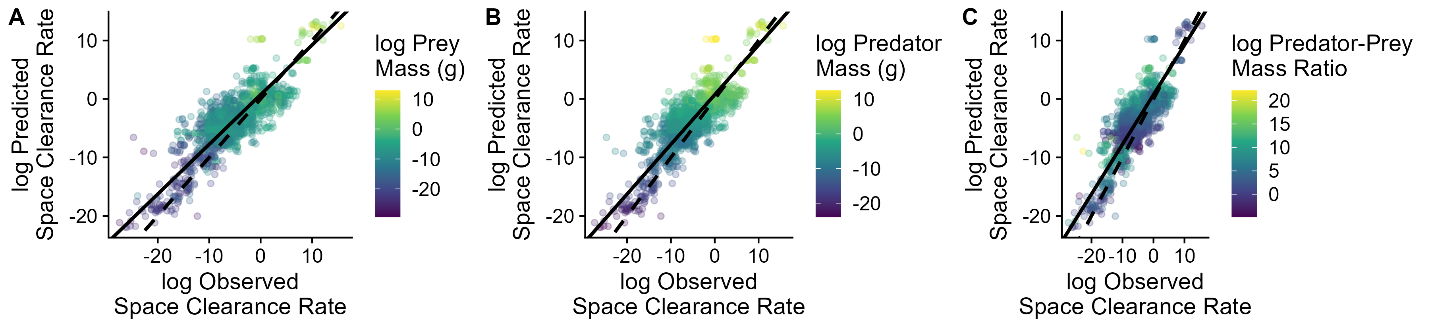

**Figure S4.1.** Observed and predicted space clearance rates from the theory colored by prey body mass (A), predator body mass (B), and the ratio of predator and prey body masses (C). Dashed lines are 1:1 lines and the solid lines are the fitted lines from a major axis regression. These plots are the same as that in Figure 1A of the main text but with body mass color coding.

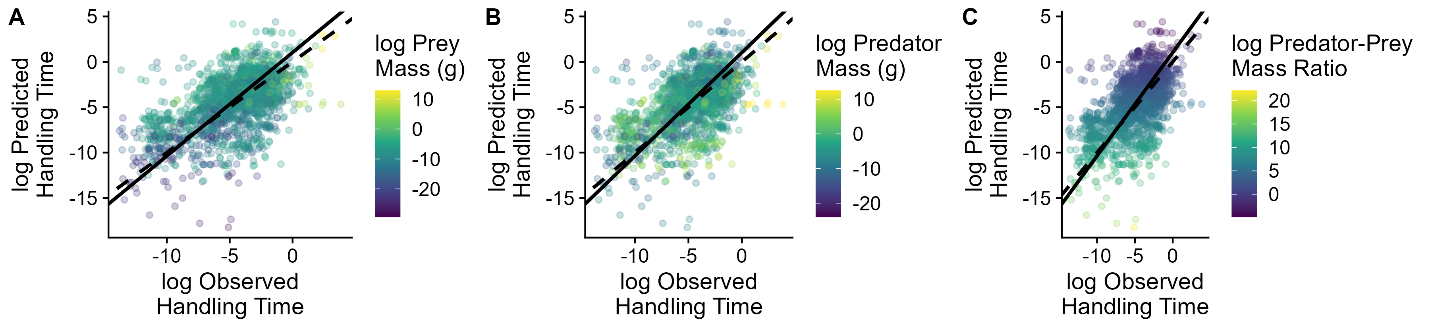

**Figure S4.2.** Observed and predicted handling times from the theory colored by prey body mass (A), predator body mass (B), and the ratio of predator and prey body masses (C). Dashed lines are 1:1 lines and the solid lines are the fitted lines from a major axis regression. These plots are the same as that in Figure 1B of the main text but with body mass color coding.

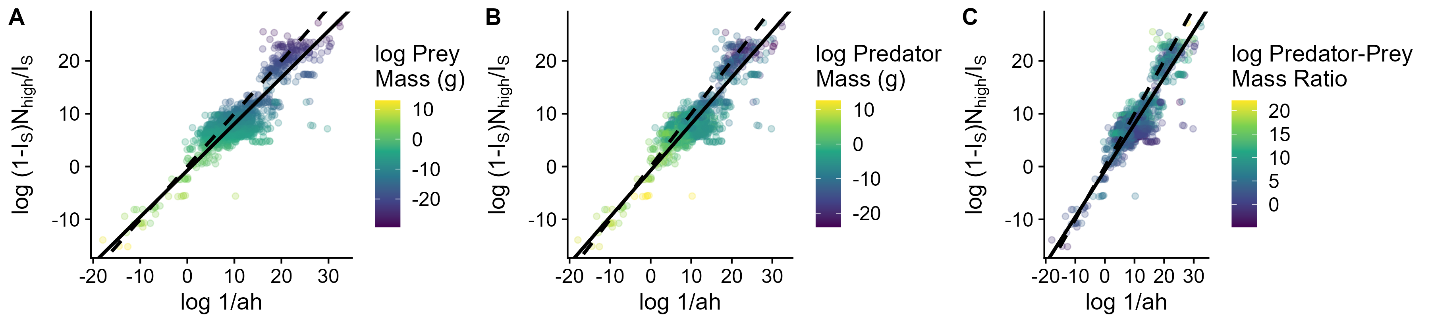

**Figure S4.3.** The theory predicts the observed half-saturation constants (*1/ah*) from the estimates of the high prey densities experienced by predators (*N_high_*) and the saturation of predator feeding rates at high prey densities (*I_S_*). Plots are color coded by prey mass (A), predator mass (B), and the predator-prey mass ratio (C). The dashed lines are 1:1 lines and the solid lines are the fits from major axis regressions. These plots are the same as that in Figure 2A of the main text but with body mass color coding.

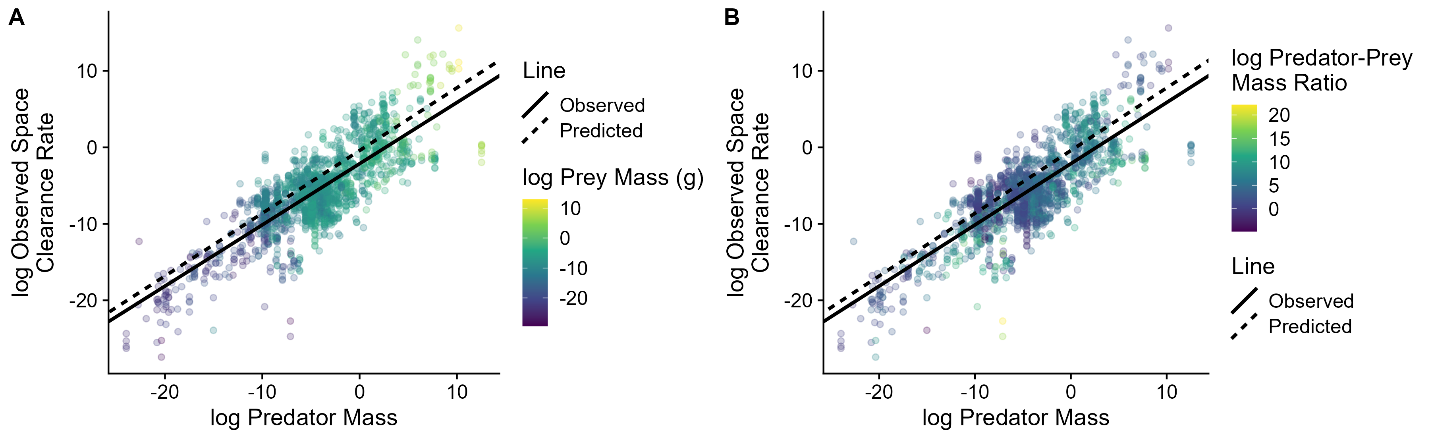

**Figure S4.4.** The theory predicts the allometric scaling of the observed space clearance rates with predator masses (in grams). Plots are color coded by prey mass (A) and predator-prey body mass ratios (B). There is no plot color coded by predator mass because this is the variable on the y-axis. The solid line is the observed allometric relationship between predator body size and space clearance rates while the dashed line is the line predicted by the theory. These plots are the same as that in Figure 3A but with the mass color coding.

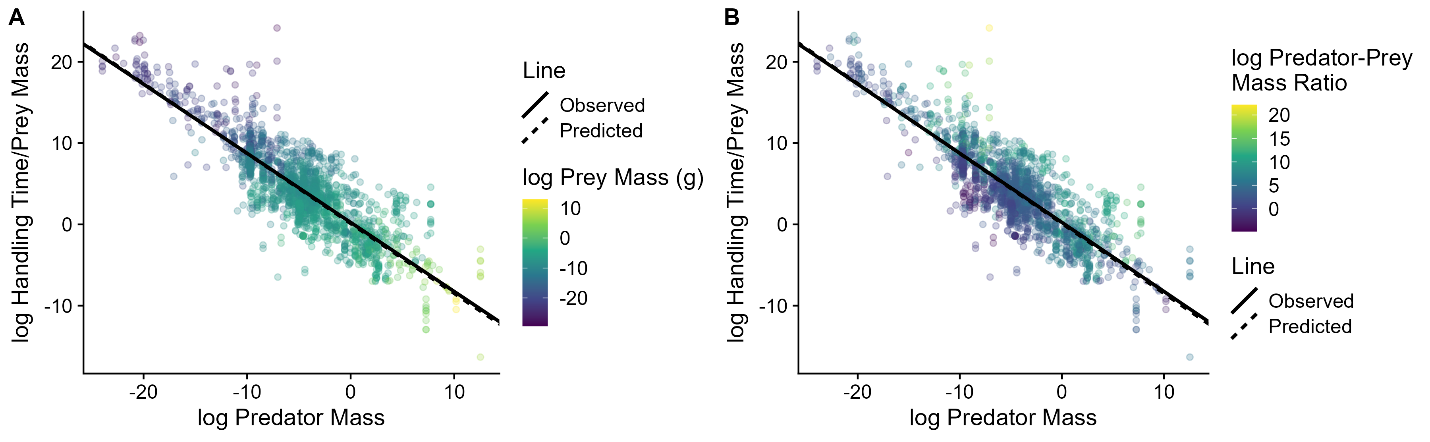

**Figure S4.5.** The theory predicts the allometric scaling of observed prey-mass-specific handling times (per gram of prey) with predator masses (in grams). Plots are color coded by prey mass (A) and predator-prey body mass ratios (B). There is no plot color coded by predator mass because this the variable on the y-axis. The solid line is the observed allometric relationship between predator body size and prey-mass-specific handling times while the dashed line the allometric relationship predicted by the theory. These plots are the same as that in Figure 3B but with the mass color coding.

**S5 Sensitivity Analysis of Choices for *I_S_*, *N_low_*, and *N_high_***

In the main text, we assume that the index of saturation at high prey densities (*I_S_*) is equal to 0.9, and that realistic low and high abundances for a prey given its mass (*N_low_* and *N_high_*) correspond to the 10^th^ and 90^th^ percentiles of the Bayesian posterior predictive distributions of empirical mass-abundance relationships. Although these are reasonable values given the rules that functional responses should meet energetic demands when prey are rare and approach their maximum when prey are abundant, there is likely to be variation among systems in the appropriate values for these parameters. For example, a particular prey may be consistently rare or abundant given its mass. Similarly, the temporal or spatial scale at which different predator populations experience variation in prey densities may alter the low and high abundances of prey they are likely to experience.

To address this, we performed sensitivity analyses to assess whether the predictive ability of the rules we propose changes given different assumptions about the values of *I_S_* and the percentiles of the predictive distribution we used to estimate *N_low_* and *N_high_*. Specifically, we examined three-way combinations of *I_S_* $\in\{$0.70, 0.9, and 0.95}, *N_low_* at the 5^th^, 10^th^ and 30^th^ percentiles of the posterior predictive distributions of abundance, and *N_high_* at the 70^th^, 90^th^, and 95^th^ percentiles of the posterior predictive distributions of abundance. We performed these sensitivity analyses for: i) the prediction of space clearance rates and handling times, i) the predicted relationship between high prey densities and the half-saturation constant, iii) the predicted relationship between space clearance rates and handling times, and iv) the allometric scaling of space clearance rates and handling times. For each of the sensitivity analyses, we reconstructed the plots and statistical analyses for the relevant combinations of *I_S_*, *N_low_*, and *N_high_*.

***Space Clearance Rate Predictions -*** Recall that the equation for the space clearance rate is $a=\frac{D}{N_{low}EM_{N}}$. Thus, for the space clearance rate, the predictions are only sensitive to the percentile used for *N_low_*. Figure S5.1 shows, for each alternative percentile of *N_low_*, the predicted versus observed values of the space clearance rates. Table S5.1 gives the correlations, *R^2^* values for the 1:1 line, and major axis regression parameters. We find little effect on our main inferences. That is, while increasing the percentile used for *N_low_* had a small positive effect on the intercept of the observed versus predicted space clearance rate relationship, it had no effect on the slope, leading to a greater underestimation of large space clearance rate values but a lower overestimation of low space clearance rate values. A potential explanation is that predators having low space clearance rates may need to meet their metabolic demands at less rare percentiles of prey abundances than predators having high space clearance rates.

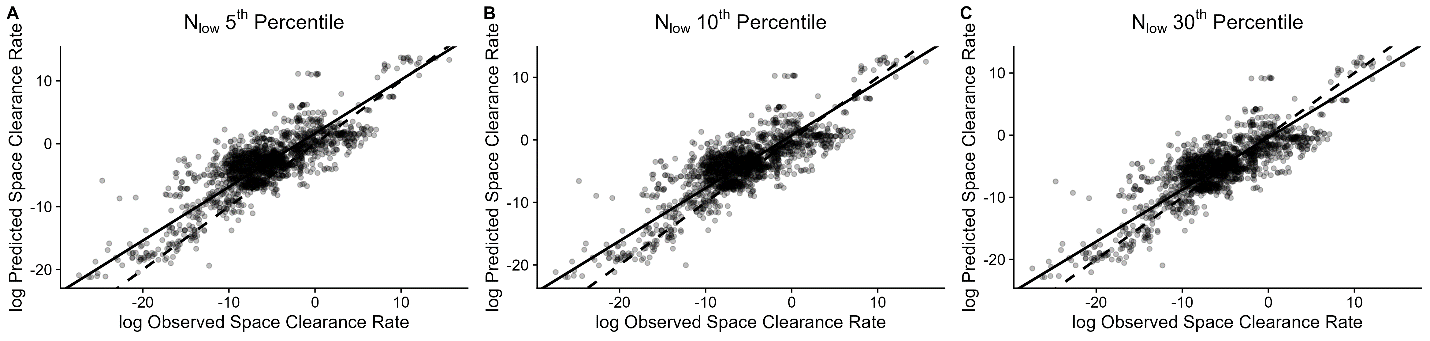

**Figure S5.1.** The relationship between observed and predicted space clearance rates is robust to changes in the percentile of the posterior predictive distribution of a mass-abundance regression used to estimate the low abundance of prey a predator is likely to experience (*N_low_*). The dashed lines are 1:1 lines and the solid lines are fits from major axis regressions.

**Table S5.1.** Summary statistics for the sensitivity analysis of space clearance rate predictions. Bolded rows highlight the combinations of parameter values used in the main text.

| ***N_low_***  **percentile** | **Correlation** | ***R*^2^**  **of 1:1** | **Intercept**  **(95% CI)** | **Slope**  **(95% CI)** |
| --- | --- | --- | --- | --- |
| 5^th^ | 0.82 | 0.75 | 1.65 (1.5,1.8) | 0.85 (0.83,0.88) |
| **10^th^** | **0.82** | **0.81** | **0.73 (0.6,0.88)** | **0.85 (0.82,0.87)** |
| 30^th^ | 0.82 | 0.85 | -0.39  (-0.53,-0.24) | 0.84 (0.81,0.86) |

***Handling Time Predictions -*** Recall that the equation for the handling time is $h=\frac{I_{S}N_{low}EM_{N}}{DN_{high}\left( 1-I_{S} \right)}$. The predictions may therefore be sensitive to variation in *N_low_*, *N_high_*, and *I_S_*. Figure S5.2 shows the predicted versus observed values of the handling time for each combination of the varied parameters. Table S5.2 gives the correlation coefficients, the *R^2^* of the 1:1 line, and major axis regression parameters. We find little effect on our main inferences, with the highest mismatch between predicted and observed handling times occurring for the combination(s) in which *I_S_* was highest and the percentile used for *N_high_* was lowest (Table S5.2).

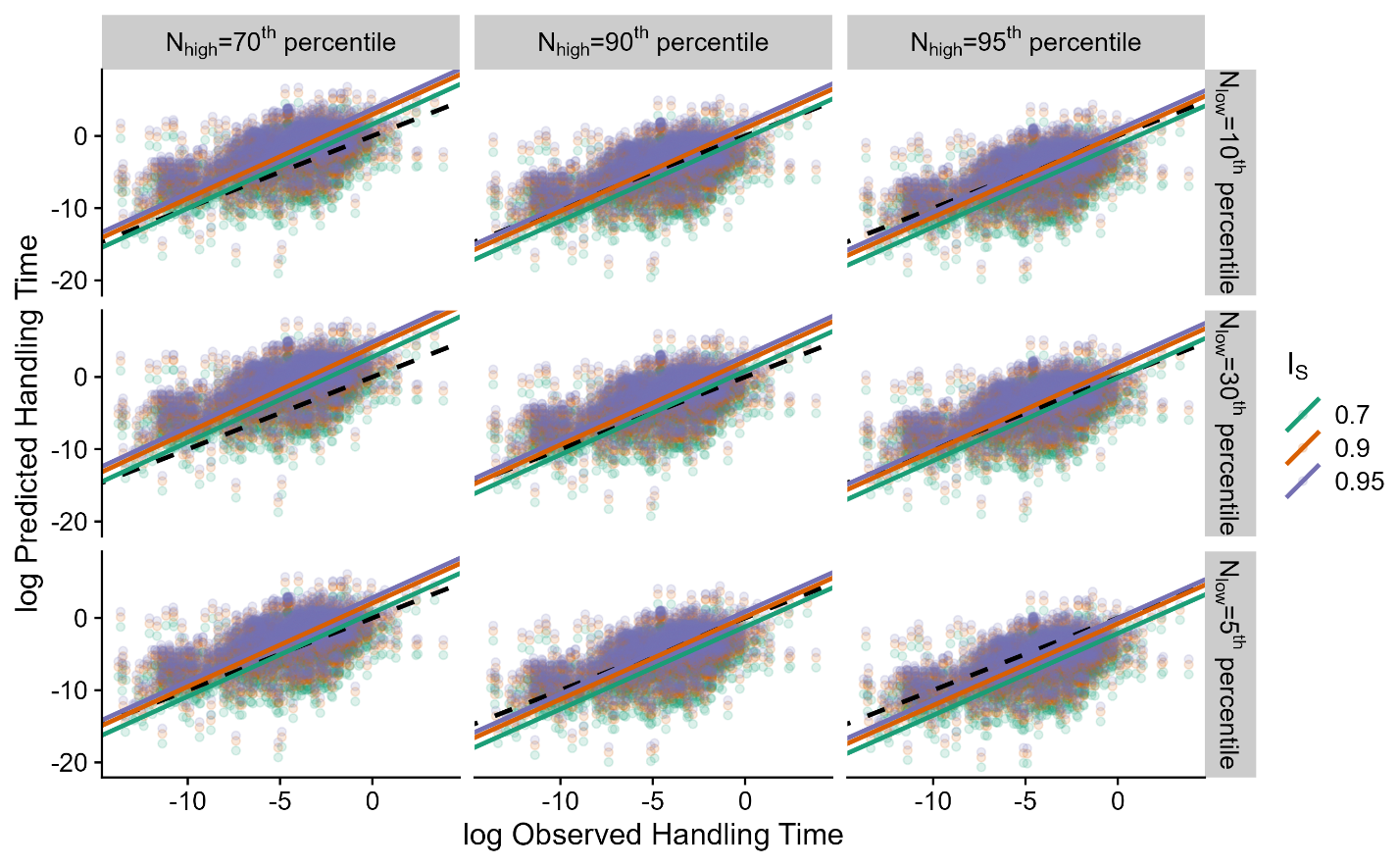

**Figure S5.2.** The relationship between observed and predicted handling times is robust to changes in the percentiles of posterior predictive distributions from mass-abundance regressions used to estimate the high prey abundances (*N_high_*) and low prey abundances (*N_low_*) that predators are likely to experience and the degree to which predator feeding rates are saturated at high prey densities (*I_S_*). The dashed lines represent the 1:1 line and the solid lines are fits from major axis regressions.

**Table S5.2.** Summary statistics for the sensitivity analysis of handling time predictions. Bolded rows highlight the combinations of parameter values used in the main text.

| ***N_low_***  **percentile** | ***N_high_***  **percentile** | ***I_S_*** | **Correlation** | ***R*^2^**  **of 1:1** | **Intercept**  **(95% CI)** | **Slope**  **(95% CI)** |
| --- | --- | --- | --- | --- | --- | --- |
| 5^th^ | 70^th^ | 0.7 | 0.574 | 0.75 | 0.70 (0.38,1.04) | 1.16  (1.09,1.23) |
| 5^th^ | 70^th^ | 0.9 | 0.574 | 0.70 | 2.05 (1.73,2.39) | 1.16  (1.09,1.23) |
| 5^th^ | 70^th^ | 0.95 | 0.574 | 0.61 | 2.80 (2.5,3.13) | 1.16  (1.09,1.23) |
| 5^th^ | 90^th^ | 0.7 | 0.571 | 0.64 | -1.22  (-1.5,-0.89) | 1.14  (1.07,1.21) |
| 5^th^ | 90^th^ | 0.9 | 0.571 | 0.74 | 0.13  (-0.19,0.46) | 1.14  (1.07,1.21) |
| 5^th^ | 90^th^ | 0.95 | 0.571 | 0.75 | 0.87  (0.56,1.21) | 1.14  (1.07,1.21) |
| 5^th^ | 95^th^ | 0.7 | 0.569 | 0.5 | -2.14  (-2.45,-1.8) | 1.14  (1.07,1.21) |
| 5^th^ | 95^th^ | 0.9 | 0.569 | 0.69 | -0.79  (-1.1,-0.46) | 1.13  (1.07,1.2) |
| 5^th^ | 95^th^ | 0.95 | 0.569 | 0.74 | -0.04  (-0.36,0.29) | 1.13  (1.07,1.2) |
| 10^th^ | 70^th^ | 0.7 | 0.575 | 0.73 | 1.62  (1.3,1.96) | 1.16  (1.1,1.24) |
| 10^th^ | 70^th^ | 0.9 | 0.575 | 0.59 | 2.97  (2.65,3.31) | 1.16  (1.1,1.24) |
| 10^th^ | 70^th^ | 0.95 | 0.575 | 0.46 | 3.72  (3.4,4.06) | 1.16  (1.1,1.24) |
| 10^th^ | 90^th^ | 0.7 | 0.573 | 0.72 | -0.31  (-0.62,0.03) | 1.15  (1.08,1.22) |
| **10^th^** | **90^th^** | **0.9** | **0.573** | **0.75** | **1.04**  **(0.73,1.38)** | **1.15**  **(1.08,1.22)** |
| 10^th^ | 90^th^ | 0.95 | 0.573 | 0.71 | 1.79  (1.47,2.12) | 1.15  (1.08,1.22) |
| 10^th^ | 95^th^ | 0.7 | 0.571 | 0.64 | -1.23  (-1.54,-0.89) | 1.14  (1.07,1.21) |
| 10^th^ | 95^th^ | 0.9 | 0.571 | 0.74 | 0.119  (-0.19,0.46) | 1.14  (1.07,1.21) |
| 10^th^ | 95^th^ | 0.95 | 0.571 | 0.75 | 0.87  (0.55,1.2) | 1.14  (1.07,1.21) |
| 30^th^ | 70^th^ | 0.7 | 0.576 | 0.63 | 2.76  (2.44,3.1) | 1.18  (1.11,1.25) |
| 30^th^ | 70^th^ | 0.9 | 0.576 | 0.40 | 4.11  (3.79,4.45) | 1.18  (1.11,1.25) |
| 30^th^ | 70^th^ | 0.95 | 0.576 | 0.21 | 4.85  (4.53,5.2) | 1.18  (1.11,1.25) |
| 30^th^ | 90^th^ | 0.7 | 0.575 | 0.75 | 0.82  (0.5,1.16) | 1.16  (1.09,1.23) |
| 30^th^ | 90^th^ | 0.9 | 0.575 | 0.68 | 2.17  (1.85,2.51) | 1.16  (1.09,1.23) |
| 30^th^ | 90^th^ | 0.95 | 0.575 | 0.6 | 2.92  (2.6,3.25) | 1.16  (1.09,1.23) |
| 30^th^ | 95^th^ | 0.7 | 0.573 | 0.73 | -0.11  (-0.42,0.23) | 1.15  (1.08,1.22) |
| 30^th^ | 95^th^ | 0.9 | 0.573 | 0.74 | 1.24  (0.93,1.58) | 1.15  (1.08,1.22) |
| 30^th^ | 95^th^ | 0.95 | 0.573 | 0.7 | 1.99  (1.68,2.33) | 1.15  (1.08,1.22) |

***Prediction of the Relationship between High Prey Densities and the Half-Saturation Constant –*** Recall that the theory predicts that the predator half-saturation constant ($\frac{1}{ah}$) is related to the high prey densities (*N_high_*) as $\frac{1}{ah}=\frac{\left( 1-I_{S} \right)N_{high}}{I_{S}}$. Since we use the observed space clearance rates and handling times from FoRAGE in our assessment of the relationship, this means that our results are potentially only sensitive to changes in the values used for *I_S_* and the posterior predictive percentiles used for *N_high_*. Figure S5.3 shows the predicted relationship between high prey densities and the half-saturation constant along with the 1:1 line for each combination of *I_S_* and the percentile of the posterior predictive distribution used to calculate *N_high_*. Table S5.3 gives the correlation coefficients and major axis regression results. We find little effect on our main inferences, with the highest mismatches in the relationship between the half-saturation constant and high prey densities occurring for the combination(s) in which *I_S_* was highest and the percentile used for *N_high_* was lowest (Figure S5.3; Table S5.3). We find small differences in correlation coefficients and slopes because changing the percentiles of the high prey densities and the value of *I_S_* merely shifts the estimates up and down.

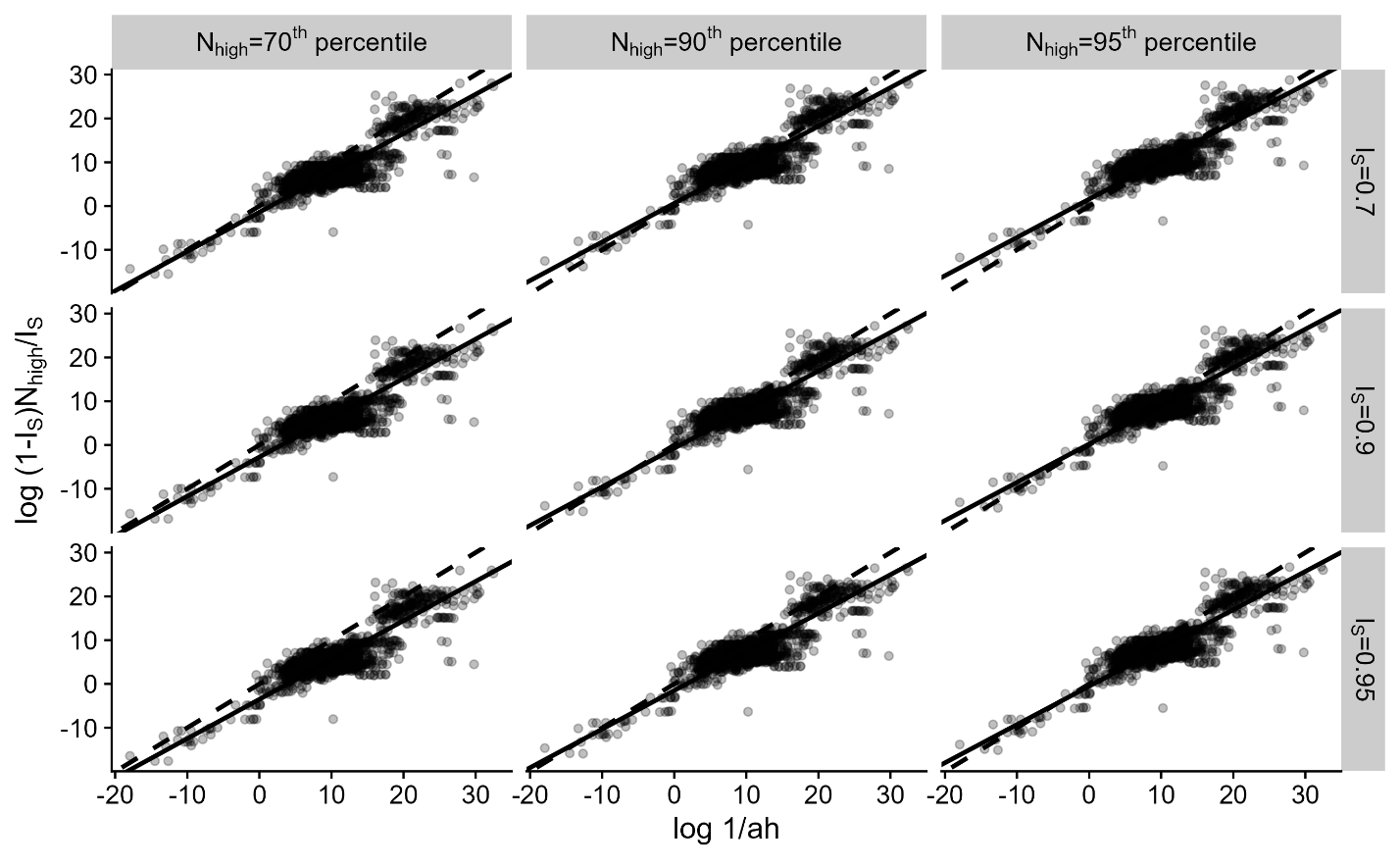

**Figure S5.3.** The predicted relationship between the half-saturation constant ($\frac{1}{ah}$) and high prey densities is largely robust to changes in the percentiles of posterior predictive distributions from mass-abundance regressions used to estimate the high prey abundances (*N_high_*) and the degree to which predator feeding rates are saturated at high prey densities (*I_S_*). The dashed lines represent the 1:1 line and the solid lines are fits from major axis regressions.

**Table S5.3.** Summary statistics for the sensitivity analysis for the relationship between the half-saturation constant and the high densities of prey. The bolded row highlights the parameter combination used in the main text.

| ***I_S_*** | ***N_high_***  **percentile** | **Correlation Coefficient** | **Intercept**  **(95% CI)** | **Slope**  **(95% CI)** |
| --- | --- | --- | --- | --- |
| 0.7 | 70^th^ | 0.86 | -1.4  (-1.64,-1.17) | 0.897  (0.875,0.92) |
| 0.7 | 90^th^ | 0.86 | 0.609  (0.373,0.84) | 0.882  (0.86,0.9) |
| 0.7 | 95^th^ | 0.86 | 1.57  (1.34,1.8) | 0.9  (0.875,0.92) |
| 0.9 | 70^th^ | 0.86 | -2.75  (-2.99,-2.52) | 0.897  (0.875,0.92) |
| **0.9** | **90^th^** | **0.86** | **-0.741**  **(-0.98,-0.51)** | **0.882**  **(0.86,0.9)** |
| 0.9 | 95^th^ | 0.86 | 0.224  (-0.01,0.45) | 0.9  (0.875,0.92) |
| 0.95 | 70^th^ | 0.86 | -3.5  (-3.74,-3.26) | 0.897  (0.875,0.92) |
| 0.95 | 90^th^ | 0.86 | -1.49  (-1.72,-1.26) | 0.882  (0.86,0.9) |
| 0.95 | 95^th^ | 0.86 | -0.52  (-0.76,-0.29) | 0.9  (0.875,0.92) |

***Prediction of the Relationship between Space Clearance Rates and Handling Times –*** Our theory predicts the relationship between space clearance rates and handling times as $ln\left( a \right)=ln\left( \frac{I_{S}}{\left( 1-I_{S} \right)N_{high}} \right)-ln\left( h \right)$. Since we use the observed space clearance rates and handling times from FoRAGE in our assessment of the relationship, this means that our results are potentially only sensitive to changes in the values used for *I_S_* and the posterior predictive percentiles used for *N_high_*. Figure S5.4 shows the predicted relationship between the handling times modified by the *I_S_* and *N_high_* and the space clearance rates along with the 1:1 line for each combination of *I_S_* and the percentile of the posterior predictive distribution used to calculate *N_high_*. Table S5.4 gives the correlation and major axis regression results. Overall, we find little sensitivity of our results to changes in *I_S_* and *N_high_*.

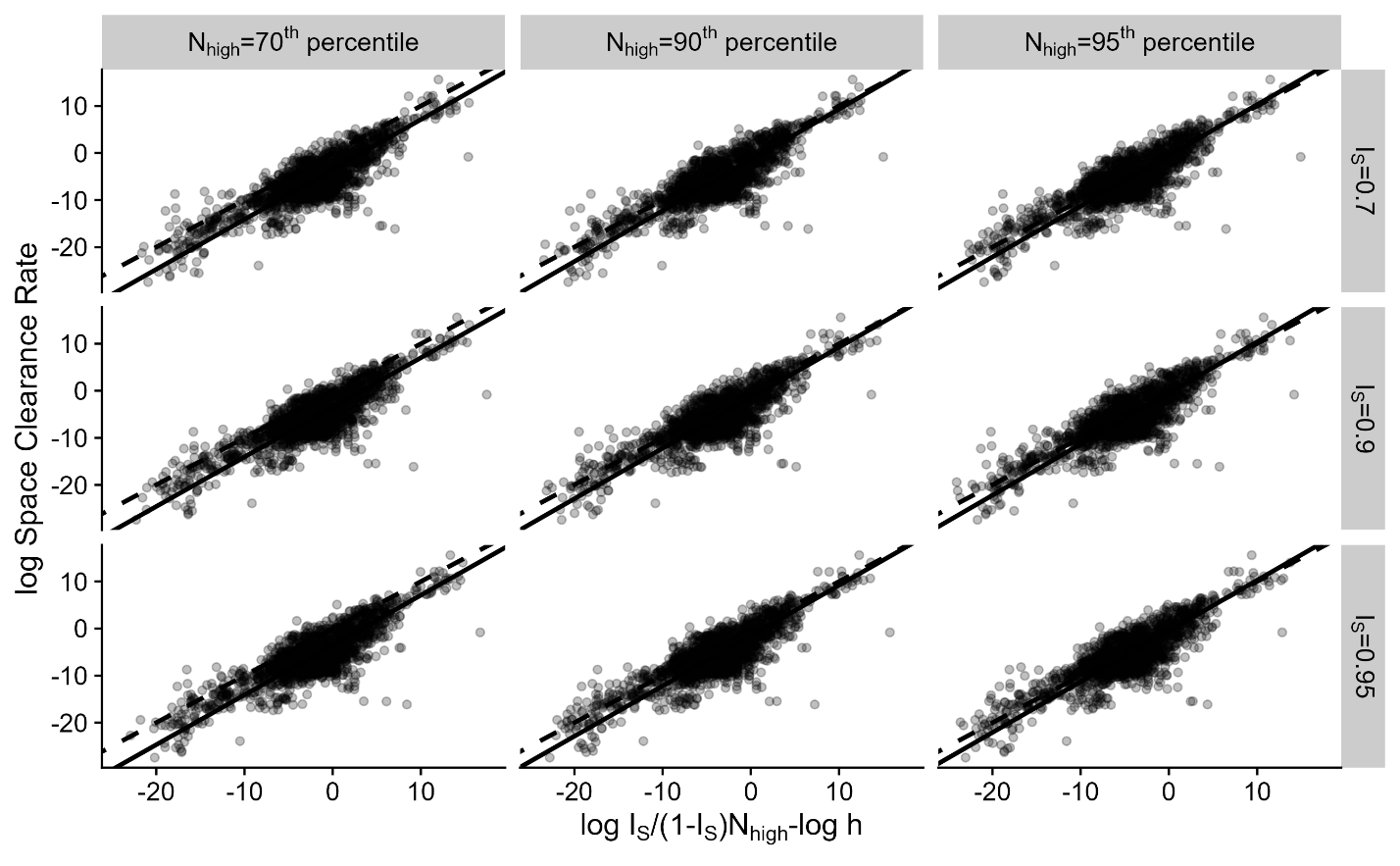

**Figure S5.4.** The predicted relationship between space clearance rates (a) and handling times (h) given high prey densities (N_high_) and the degree of saturation in predator feeding rates (I_S_) is largely robust to changes in the percentiles of posterior predictive distributions from mass-abundance regressions used to estimate the high prey abundances and the values of *I_S_*. The dashed lines represent the 1:1 line and the solid lines are fits from major axis regressions.

**Table S5.4.** Summary statistics for the sensitivity analysis for the relationship between space clearance rates and handling times. The bolded row highlights the parameter combination used in the main text.

| ***I_S_*** | ***N_high_***  **percentile** | **Correlation Coefficient** | **Intercept**  **(95% CI)** | **Slope**  **(95% CI)** |
| --- | --- | --- | --- | --- |
| 0.7 | 70^th^ | 0.836 | -3.49  (-3.56,-3.43) | 1.06  (1.03,1.09) |
| 0.7 | 90^th^ | 0.837 | 1.46  (-1.58,-1.34) | 1.08  (1.05,1.11) |
| 0.7 | 95^th^ | 0.835 | -0.51  (-0.66,-0.36) | 1.08  (1.05,1.11) |
| 0.9 | 70^th^ | 0.837 | -3.51  (-3.57,-3.45) | 1.05  (1.02,1.08) |
| **0.9** | **90^th^** | **0.836** | **-1.49**  **(-1.61,-1.37)** | **1.07**  **(1.04,1.1)** |
| 0.9 | 95^th^ | 0.836 | -0.47  (-0.62,-0.32) | 1.08  (1.06,1.12) |
| 0.95 | 70^th^ | 0.836 | -3.5  (-3.56,-3.43) | 1.06  (1.03,1.09) |
| 0.95 | 90^th^ | 0.837 | -1.51  (-1.62,-1.39) | 1.07  (1.04,1.1) |
| 0.95 | 95^th^ | 0.838 | -0.51  (-0.66,-0.37) | 1.08  (1.05,1.11) |

***Prediction of the Allometric Scaling of Space Clearance Rates –*** Equation S2.3 shows that space clearance rates are expected to scale with predator body mass as

$a= \frac{\delta_{0}\mu_{0,P}^{-\eta_{low}-1}}{\eta_{0,low}E}M_{P}^{\delta+\mu_{P}\left( -\eta_{low}-1 \right)}$.

The definitions of the parameters are given in table S2.1. This equation shows that, among the variables for which we examine the sensitivity of our predictions, the predicted allometric scaling is only sensitive to the slope and intercept of the relationship between prey body mass and the low density of prey ($\eta_{0,\mathrm{low}}$ and $\eta_{low}$). Therefore, we examined the sensitivity of our allometric predictions by estimating the intercept and slope of the relationship between prey body masses and low prey densities estimated at the 5^th^, 10^th^, and 30^th^ percentiles of the posterior predictive distribution of the relationship between prey mass and population density. We then used these intercepts and slopes to calculate the predicted intercept and slope of the allometric scaling between predator mass and space clearance rates with all of the other parameters at the values given in table S2.1.

Overall, we found little variation in the predicted slope of the allometric relationship between space clearance rates and predator masses, with all slopes being within the confidence interval of the estimate of the observed slope (estimated value = 0.8; 95% confidence interval (CI) = 0.78, 0.83; Table S5.4, Figure S5.4). The predicted intercepts showed greater variation, with the low prey densities estimated at the 30^th^ percentile having the intercept closest to the intercept of the observed allometric relationship (estimated value = -2.14; 95% CI = -2.3,-1.97). We conclude that our results for the prediction of the allometric relationship between predator masses and space clearance rates are robust to the assumptions made in the main text.

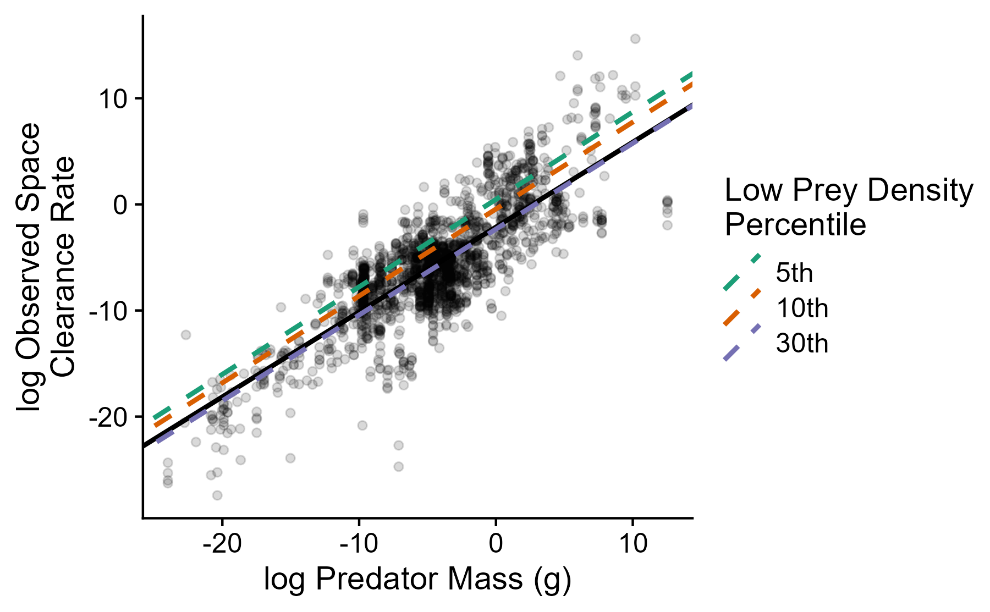

**Figure S5.5.** Predicted allometric relationships between predator masses and space clearance rates are largely robust to the use of different percentiles to estimate the scaling relationship between prey masses and their low densities. The solid line is the observed allometric relationship between predator masses and space clearance rates.

**Table S5.5.** Summary statistics for the sensitivity analysis for the allometric relationship between predator masses and space clearance rates. The bolded row highlights the parameter combination used in the main text.

| ***N_low_***  **percentile** | $\boldsymbol{\eta}_{\boldsymbol{0,low}}$ | $\boldsymbol{\eta}_{\boldsymbol{low}}$ | **Predicted Allometric Intercept** | **Predicted Allometric Slope** |
| --- | --- | --- | --- | --- |
| 5^th^ | 0.009 | -0.95 | 0.47 | 0.82 |
| **10^th^** | **0.02** | **-0.95** | **-0.44** | **0.82** |
| 30^th^ | 0.16 | -0.93 | -2.33 | 0.81 |

***Prediction of the Allometric Scaling of Handling Times –*** Equation S2.4 shows that handling times are predicted to scale with predator and prey body masses as

$h=\frac{I_{S}\eta_{0,low}E}{\left( 1-I_{S} \right)\delta_{0}\eta_{0,high}}M_{N}^{1+\eta_{low}-\eta_{high}}M_{P}^{-\delta}$.

The definitions of all parameters are given in Table S2.1. This equation shows that both the prefactor and the scaling relationship between handling times and masses may be sensitive to the variables for which we examine the sensitivity of our predictions. In S2, we further consider the approximation of equation S2.4 as

$\frac{h}{M_{N}}=\frac{I_{S}\eta_{0,low}E}{\left( 1-I_{S} \right)\delta_{0}\eta_{0,high}}M_{P}^{-\delta}$.

This is because the values for $\eta_{low}$ and $\eta_{high}$ were such that the exponent for $M_{N}$ was approximately 1.

Allowing the percentiles used to determine the relationship between prey masses and high and low prey abundances to vary, we again obtain values for the exponent of $M_{N}$ that ranged from 0.94-0.97 across all the considered combinations of $\eta_{low}$ and $\eta_{high}$ values estimated at the different percentiles. We therefore conclude that this approximation is valid regardless of the percentiles used. As the scaling exponent then only depends on $\delta$ (the slope of the allometric relationship between predator mass and metabolic rate), the slope of the allometric relationship prey-mass-specific handling times and predator masses is not sensitive to changes in any of the variables that we vary (Figure S5.5). The predicted intercepts of the allometric scaling relationship between prey-mass-specific handling times and predator mass do vary, although the predicted values, in general, do not fall far from the observed intercept value of 0.24 (95% CI = 0.08,0.4; Figure S5.5, Table S5.5). We conclude that our results are generally robust to reasonable choices in the percentiles used to determine the relationship between high and low prey densities and prey masses and the values for *I_S_*.

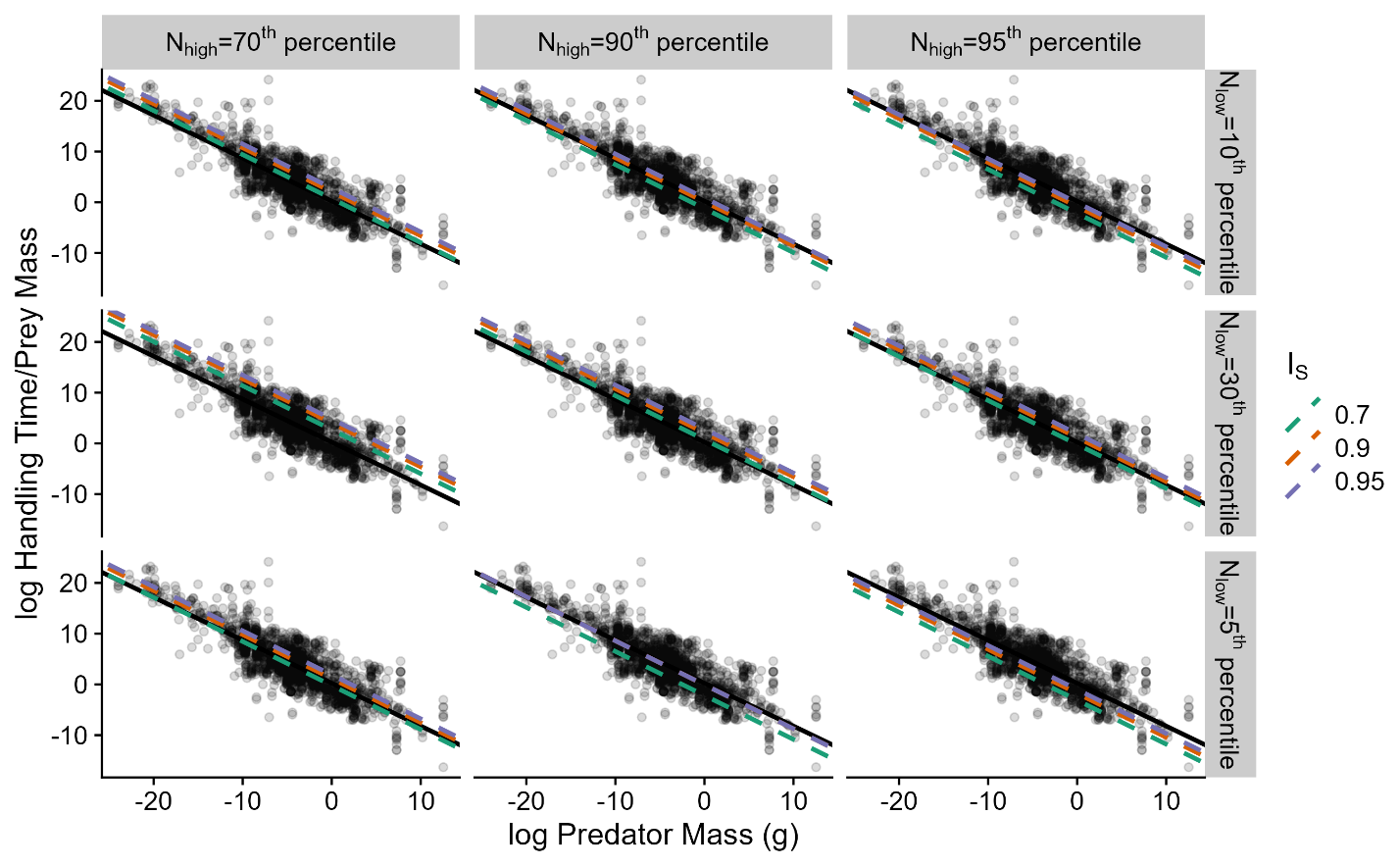

**Figure S5.6.** Predicted allometric relationships between predator masses and prey-mass-specific handling times (per gram of prey) are largely robust to the use of different percentiles to estimate the scaling relationship between prey masses and their low and densities and the degree of predator feeding rate saturation at high prey densities (*I_S_*). The solid line is the observed allometric relationship between predator masses and space clearance rates.

**Table S5.6.** Summary statistics for the sensitivity analysis for the allometric relationship between predator masses and prey-mass-specific handling times. The bolded row highlights the parameter combination used in the main text.

| ***I_S_*** | ***N_high_* percentile** | ***N_low_***  **percentile** | $\boldsymbol{\eta}_{\boldsymbol{0,low}}$ | $\boldsymbol{\eta}_{\boldsymbol{low}}$ | $\boldsymbol{\eta}_{\boldsymbol{0,high}}$ | $\boldsymbol{\eta}_{\boldsymbol{high}}$ | **Predicted Allometric Intercept** |
| --- | --- | --- | --- | --- | --- | --- | --- |
| 0.7 | 70^th^ | 5^th^ | 0.009 | -0.95 | 2.42 | -0.91 | -0.22 |
| 0.7 | 70^th^ | 10^th^ | 0.02 | -0.95 | 2.42 | -0.91 | 0.72 |
| 0.7 | 70^th^ | 30^th^ | 0.16 | -0.93 | 2.42 | -0.91 | 2.68 |
| 0.7 | 90^th^ | 5^th^ | 0.009 | -0.95 | 17.15 | -0.9 | -2.18 |
| 0.7 | 90^th^ | 10^th^ | 0.02 | -0.95 | 17.15 | -0.9 | -1.24 |
| 0.7 | 90^th^ | 30^th^ | 0.16 | -0.93 | 17.15 | -0.9 | 0.72 |
| 0.7 | 95^th^ | 5^th^ | 0.009 | -0.95 | 43.86 | -0.89 | -3.12 |
| 0.7 | 95^th^ | 10^th^ | 0.02 | -0.95 | 43.86 | -0.89 | -2.18 |
| 0.7 | 95^th^ | 30^th^ | 0.16 | -0.93 | 43.86 | -0.89 | -0.22 |
| 0.9 | 70^th^ | 5^th^ | 0.009 | -0.95 | 2.42 | -0.91 | 1.13 |
| 0.9 | 70^th^ | 10^th^ | 0.02 | -0.95 | 2.42 | -0.91 | 2.07 |
| 0.9 | 70^th^ | 30^th^ | 0.16 | -0.93 | 2.42 | -0.91 | 4.03 |
| 0.9 | 90^th^ | 5^th^ | 0.009 | -0.95 | 17.15 | -0.9 | -0.08 |
| **0.9** | **90^th^** | **10^th^** | **0.02** | **-0.95** | **17.15** | **-0.9** | **0.11** |
| 0.9 | 90^th^ | 30^th^ | 0.16 | -0.93 | 17.15 | -0.9 | 2.07 |
| 0.9 | 95^th^ | 5^th^ | 0.009 | -0.95 | 43.86 | -0.89 | -1.77 |
| 0.9 | 95^th^ | 10^th^ | 0.02 | -0.95 | 43.86 | -0.89 | -0.83 |
| 0.9 | 95^th^ | 30^th^ | 0.16 | -0.93 | 43.86 | -0.89 | 1.13 |
| 0.95 | 70^th^ | 5^th^ | 0.009 | -0.95 | 2.42 | -0.91 | 1.88 |
| 0.95 | 70^th^ | 10^th^ | 0.02 | -0.95 | 2.42 | -0.91 | 2.82 |
| 0.95 | 70^th^ | 30^th^ | 0.16 | -0.93 | 2.42 | -0.91 | 4.78 |
| 0.95 | 90^th^ | 5^th^ | 0.009 | -0.95 | 17.15 | -0.9 | -0.08 |
| 0.95 | 90^th^ | 10^th^ | 0.02 | -0.95 | 17.15 | -0.9 | 0.86 |
| 0.95 | 90^th^ | 30^th^ | 0.16 | -0.93 | 17.15 | -0.9 | 2.82 |
| 0.95 | 95^th^ | 5^th^ | 0.009 | -0.95 | 43.86 | -0.89 | -1.02 |
| 0.95 | 95^th^ | 10^th^ | 0.02 | -0.95 | 43.86 | -0.89 | -0.08 |
| 0.95 | 95^th^ | 30^th^ | 0.16 | -0.93 | 43.86 | -0.89 | 1.88 |

**S6 Comparison of Allometric Scaling Predictions with Yodzis and Innes (1992)**

In the Discussion of the main text, we compare the allometric scaling of the functional response parameters that are observed in FoRAGE and predicted by our theory to the allometric scaling relationships in previous studies. Here, we derive our comparison between the allometric relationships we observed and those derived from Yodzis and Innes (1992; hereafter Y&I), one of the first papers introducing empirical allometric scaling relationships into consumer-resource models.

Making the comparison between the allometric relationships observed and derived in this manuscript to those in Y&I is more complex than it might seem at face value. This is because Y&I consider the functional response in different units and use the Michaelis-Menten version of the Type II functional response rather than the Holling Disc Equation version that we use (Holling 1959a). Below, we walk through the steps required to compare the allometric scaling relationships and show that the scaling implied by Y&I is quite close to the scaling that we derive and observe.

***Scaling in Yodzis and Innes (1992)***

Y&I examine the dynamics of consumer and resource biomass and assume that the functional response of the consumer is

$J\left( R \right)=\frac{J_{max}R}{R_{0}+R}$	eqn. S6.1

where $J\left( R \right)$ is the per consumer mass feeding rate of the consumer on the resource, $J_{max}$ is the maximum per consumer mass feeding rate, $R$ is the resource mass density, and $R_{0}$is the half-saturation constant in prey mass. They argue that $J_{max}$ scales with predator mass ($M_{P}$) to the same exponent as metabolic rates with predator mass which we will call $\mu$. Because Y&I $J_{max}$ is predator-mass specific, $J_{max}$ should scale to an exponent of $\mu- 1$ (because $\frac{M_{P}^{\mu}}{M_{P}}=M_{P}^{\mu-1}$). Although Y&I do not explain their reasoning in their text, they consider $R_{0}$ to be independent of predator and prey masses.

***Comparing the scaling of handling time to the scaling of*** $\boldsymbol{J}_{\boldsymbol{max}}$

We begin comparing the allometric scaling relationships in our theory and the FoRAGE database to those of Y&I by comparing the allometric scaling relationships of handling time ($h$) in our manuscript to the allometric scaling of $J_{max}$ in Y&I. Again, Y&I state that $J_{max}\propto M_{P}^{\mu-1}$. In our manuscript, we find that the prey mass-specific handling time scales with predator mass as $\frac{h}{M_{N}} \propto M_{P}^{\mu}$ (See Supplemental Material S2).

To compare the allometric scaling relationships we need to convert from one to the other such that they are in the same units. To begin we will consider a parameter $\eta$ which will be the handling time of the Holling Disc Equation form of the Type II functional response in the units of Y&I. We know that the conversion between the maximum feeding rate, $J_{max}$, in the Michaelis-Menten version of the Type II functional response to the handling time, $\eta$ is

$\frac{1}{\eta}=J_{max}$.	eqn. S6.2

We also know that $J_{max}$ has units $\frac{\left[ \text{prey mass} \right]}{\left[ \text{time} \right]\left[ \text{pred mass} \right]}$. Therefore, $\eta$ has units $\frac{\left[ \text{time} \right]\left[ \text{pred mass} \right]}{\left[ \text{prey mass} \right]}$. Our $h$ is in units of predator and prey individuals, so is $\frac{\left[ \text{time} \right]\left[ \text{predators} \right]}{\left[ \text{prey} \right]}$. To get $h$ into the same units $\eta$ then, we need to be able to convert between predator and prey numbers and masses. To do so, we assume that the mass we assign to a predator or prey is the representative average mass of an individual in the population. In the case, if $N$ is the number of individuals in the population and $B_{N}$ is the biomass of the population, then $N=B_{N}/M_{N}$. Hence, we can convert from numbers to masses by multiplying the numbers by the average mass of an individual in the population ($NM_{N}=B_{N}$).

Now that we know how to translate between $J_{max}$ and $h$, we can compare the allometric scaling relationship that we derive to that assumed by Y&I. We have that

$\frac{h}{M_{N}}\propto M_{P}^{-\mu}$	eqn. S6.3

The left-hand side of this equation has units $\frac{\left[ \text{time} \right]\left[ \text{predators} \right]}{\left[ \text{prey mass} \right]}$. To convert this to $\eta$, we need to multiply by predator mass. This gives us

$\frac{hM_{P}}{M_{N}}=\eta\propto M_{P}^{-\mu+1}$.	eqn. S6.4

Next, we take the reciprocal of both sides to get

$\frac{1}{\eta}=J_{max}\propto M_{P}^{\mu-1}$.	eqn. S6.5

Therefore, we see that the allometric relationship that we derive for the handling time is equivalent to the allometric relationship implied by Y&I that the predator mass-specific maximum feeding rate in terms of prey mass scales in the same was as the predator’s mass-specific metabolic rate.

***Comparing the scaling of space clearance rate to that implied by Y&I***

Now we will examine the implications of Y&I’s allometric scaling for the space clearance rate and compare that to our derived allometric scaling relationship. Again, Y&I assume that $J_{max}\propto M_{P}^{\mu-1}$ and that $R_{0}$ (the half-saturation constant in units of prey mass) has no relationship with predator or prey masses. If we let $\alpha$ and $\eta$ be the space clearance rate and handling times of the Holling disc equation in the units of Y&I, we can again convert between the Michaelis-Menten and Holling forms of the Type II functional response as

$\frac{1}{\alpha\eta}=R_{0}$.

Next, we isolate $\alpha$.

$\frac{1}{\alpha}=\eta R_{0}$	eqn. S6.6

$\alpha=\frac{1}{\eta R_{0}}$	eqn. S6.7

Now, using that $\frac{1}{\eta}=J_{max}$, we finally get

$\alpha=\frac{J_{max}}{R_{0}}$	eqn. S6.8

Because $R_{0}$ is assumed not to scale with prey or predator densities and $J_{max}\propto M_{P}^{\mu-1}$, $\alpha$ should also scale with predator mass to the $\mu- 1$ power. Now to convert from the scaling of $\alpha$ to our scaling of the space clearance rate $a$ we need to consider the units of the two parameters. As implied by the equation S6.8 above along with the units of $J_{max}$ being $\frac{\left[ \text{prey mass} \right]}{\left[ \text{time} \right]\left[ \text{pred mass} \right]}$ and the units of $R_{0}$ being $\frac{\left[ \text{prey mass} \right]}{\left[ \text{space} \right]}$, the units of $\alpha$ are $\frac{\left[ \text{space} \right]}{\left[ \text{pred mass} \right]\left[ \text{time} \right]}$. The units of our space clearance rate $a$ are $\frac{\left[ \text{space} \right]}{\left[ \text{predators][}\text{time} \right]}$ which we find scales with predator mass in the FoRAGE database as $a\propto M_{P}^{0.8}$ and in our theory as $a\propto M_{P}^{0.82}$. To compare this to the allometric scaling of $\alpha$, we convert $a$ to $\alpha$ by dividing by predator mass. From our theory, we then get that $\frac{a}{M^{P}}=\alpha\propto M_{P}^{-0.18}$. Since, in our dataset, $\mu= 0.87$ (See the main text and Supplementary Material S2), Y&I’s allometric scaling implies that $\alpha\propto M_{P}^{0.13}$, which is not far from our results.

**Supplementary References**

Buckner, C.H. (1964). Metabolism, food capacity, and feeding behavior in four species of shrews. *Can. J. Zool.*, 42, 259–279.

Carbone, C., Teacher, A. & Rowcliffe, J.M. (2007). The Costs of Carnivory. *PLOS Biology*, 5, e22.

Chan, K., Boutin, S., Hossie, T.J., Krebs, C.J., O’Donoghue, M. & Murray, D.L. (2017). Improving the assessment of predator functional responses by considering alternate prey and predator interactions. *Ecology*, 98, 1787–1796.

Chizzolini, R., Zanardi, E., Dorigoni, V. & Ghidini, S. (1999). Calorific value and cholesterol content of normal and low-fat meat and meat products. *Trends in Food Science & Technology*, 10, 119–128.

Coblentz, K.E., Novak, M. & DeLong, J.P. (2023). Predator feeding rates may often be unsaturated under typical prey densities. *Ecology Letters*, 26, 302–312.

Cook, J.G., Kelly, A.P., Cook, R.C., Culling, B., Culling, D., McLaren, A., *et al.* (2021). Seasonal patterns in nutritional condition of caribou (Rangifer tarandus) in the southern Northwest Territories and northeastern British Columbia, Canada. *Can. J. Zool.*, 99, 845–858.

Dale, B.W., Adams, L.G. & Bowyer, R.T. (1994). Functional Response of Wolves Preying on Barren-Ground Caribou in a Multiple-Prey Ecosystem. *Journal of Animal Ecology*, 63, 644–652.

Gorecki, A. (1965). Energy values of body in small mammals. *Acta Theriologica*, 10, 333–352.

Holling, C.S. (1959a). Some Characteristics of Simple Types of Predation and Parasitism. *Can Entomol*, 91, 385–398.

Holling, C.S. (1959b). The Components of Predation as Revealed by a Study of Small-Mammal Predation of the European Pine Sawfly1. *The Canadian Entomologist*, 91, 293–320.

Korpimaki, E. & Norrdahl, K. (1991). Numerical and Functional Responses of Kestrels, Short-Eared Owls, and Long-Eared Owls to Vole Densities. *Ecology*, 72, 814–826.

Menzies, A.K., Studd, E.K., Seguin, J.L., Derbyshire, R.E., Murray, D.L., Boutin, S., *et al.* (2022). Activity, heart rate, and energy expenditure of a cold-climate mesocarnivore, the Canada lynx (Lynx canadensis). *Can. J. Zool.*, 100, 261–272.

Messier, F. (1994). Ungulate Population Models with Predation: A Case Study with the North American Moose. *Ecology*, 75, 478–488.

Novak, M., Wolf, C., Coblentz, K.E. & Shepard, I.D. (2017). Quantifying predator dependence in the functional response of generalist predators. *Ecology Letters*, 20, 761–769.

O’Donoghue, M., Boutin, S., Krebs, C.J., Zuleta, G., Murray, D.L. & Hofer, E.J. (1998). Functional Responses of Coyotes and Lynx to the Snowshoe Hare Cycle. *Ecology*, 79, 1193–1208.

Schwartz, C.C., Hubbert, M.E. & Franzmann, A.W. (1988). CHANGES IN BODY COMPOSITION OF MOOSE DURING WINTER. *Alces: A Journal Devoted to the Biology and Management of Moose*, 24, 178–187.

Sundell, J., Norrdahl, K., Korpimäki, E. & Hanski, I. (2000). Functional response of the least weasel, Mustela nivalis nivalis. *Oikos*, 90, 501–508.

Uiterwaal, S.F., Lagerstrom, I.T., Lyon, S.R. & DeLong, J.P. (2022). FoRAGE database: A compilation of functional responses for consumers and parasitoids. *Ecology*, 103, e3706.

Vucetich, J.A., Peterson, R.O. & Schaefer, C.L. (2002). The Effect of Prey and Predator Densities on Wolf Predation. *Ecology*, 83, 3003–3013.

Yodzis, P. & Innes, S. (1992). Body Size and Consumer-Resource Dynamics. *The American Naturalist*, 139, 1151–1175.

Zalewski, A., Jędrzejewski, W. & Jędrzejewska, B. (1995). Pine marten home ranges, numbers and predation on vertebrates in a deciduous forest (Białowieża National Park, Poland). *Annales Zoologici Fennici*, 32, 131–144.
